# A Model-Based Approach for Pulse Selection from Electrodermal Activity

**DOI:** 10.1101/2020.05.17.098129

**Authors:** Sandya Subramanian, Patrick L. Purdon, Riccardo Barbieri, Emery N. Brown

**Affiliations:** Harvard-MIT Division of Health Sciences and Technology, Massachusetts Institute of Technology, Cambridge, MA 02139 USA; MGH Department of Anesthesia, Critical Care, and Pain Medicine, Picower Institute for Learning and Memory, and Harvard Medical School; Department of Electronics, Informatics and Engineering, Politecnico di Milano, Milano, Italy. He is also with MGH and Massachusetts Institute of Technology, MA; MGH Department of Anesthesia, Critical Care, and Pain Medicine, MIT Department of Brain and Cognitive Sciences, MIT Institute for Medical Engineering and Science, and Picower Institute for Learning and Memory, Cambridge, MA

**Keywords:** electrodermal activity, inverse Gaussian, point processes, signal processing, statistics

## Abstract

**Objective:** The goal of this work was to develop a physiology-based paradigm for pulse selection from electrodermal activity (EDA) data.

**Methods:** We aimed to use insight about the integrate-and-fire physiology of sweat gland bursts, which predicts inverse Gaussian inter-pulse interval structure. At the core of our paradigm is a subject-specific amplitude threshold selection process for pulses based on the statistical properties of four right-skewed models including the inverse Gaussian. These four models differ in their tail behavior, which reflects sweat gland physiology to varying degrees. By screening across thresholds and fitting all four models, we selected for heavier tails that reflect inverse Gaussian-like structure and verified the pulse selection with a goodness-of-fit analysis.

**Results:** We tested our paradigm on two different subject cohorts recorded during different experimental conditions and using different equipment. In both cohorts, our method robustly and consistently recovered pulses that captured the inverse Gaussian-like structure predicted by physiology, despite large differences in noise level of the data. In contrast, an established EDA analysis paradigm, which assumes a constant amplitude threshold across all data, was unable to separate pulses from noise.

**Conclusion:** We present a computationally efficient, statistically rigorous, and physiology-informed paradigm for pulse selection from EDA data that is robust across individuals and experimental conditions yet adaptable to changes in noise level.

**Significance:** The robustness of our paradigm and its basis in physiology move EDA closer to serving as a clinical marker for sympathetic activity in diverse conditions such as pain, anxiety, depression, and sleep.

## I. Introduction

Sweat gland activity is one of the most primitive parts of our “fight or flight” response, and is therefore used as a way to assess sympathetic nervous system activity in the body [1]. Sympathetic activation is induced by emotional and physiologic states such as stress, anxiety, and pain. Electrodermal activity (EDA) measures the electrical conductance of the skin as a proxy for the activity of sweat glands. As sweat glands are more active, due to physiologic or emotional stimulation, the electrical conductance of the skin increases since sweat conducts electricity [1]. The continuous measurement of EDA can be performed inexpensively and non-invasively. For this reason, it has been employed in lie detector tests and as a neuromarketing tool. There is growing interest in the development of real-time algorithms to accurately characterize EDA to track changes in emotional and physiologic states.

EDA consists of two simultaneous levels of activity. The baseline or tonic component represents background or ambient conditions and drifts gradually over minutes. On top of that, pulsatile sweat release events make up what is known as the phasic component of EDA, which has a timescale of a few seconds and is thought to correspond to sympathetic nervous system activity [1].

We have previously shown that the inter-pulse interval distribution in EDA data follows an inverse Gaussian distribution [2][3], which agrees with a model of the rise of sweat through the gland to the skin surface as an integrate-and-fire process [4][5]. This process is similar to the mechanism that underlies other point process events such as neuronal and cardiac action potentials, earthquakes, geysers, and volcanoes [6][7]. We showed that the temporal structure in EDA favors right-skewed heavy tailed distributions, including the inverse Gaussian and even heavier tailed distributions such as the lognormal, due to the presence of sparse regions of EDA with low activity and long inter-pulse intervals [2]. This result makes possible the use of low-order models in EDA analyses and increases the signal-to-noise ratio.

Nevertheless, we observed a high degree of intersubject variability in the number of pulsatile events (pulses) despite all subjects being at rest for the same duration of time [2]. This is due to the inherent variability in background filling levels and spontaneous activity of sweat glands between people, as well as differing levels of noise inherent to different recording equipment and experimental conditions. We previously assumed a relatively stable background level, but this cannot be assumed across different experimental conditions. Robust and accurate capture of the statistical structure we previously described depends critically on extracting pulses in a consistent and reliable way.

To this end, the relationship between the properties of several right-skewed distributions helps identify the presence of noise in any subject cohort. For example, while the heavier-tailed inverse Gaussian and lognormal distributions are indicators of desired statistical structure, the flexibility of the gamma distribution in capturing the light tails likely to occur with random noise in a subject cohort allows it to function as a noise indicator [2]. By examining the patterns of goodness-of-fit of four distributions (inverse Gaussian, lognormal, gamma, exponential) across subjects in a cohort, we gauged the level of noise in the subject cohort and the set of pulses to extract. We also identified individual subjects who were clear outliers from the rest of the cohort.

In this work, we define a robust method for extracting pulses from any observational EDA subject cohort and show that it is successful in capturing the underlying statistical structure in two subject cohorts collected from different populations, under different experimental conditions, and using different equipment. The method combines the steps of extracting pulses and identifying the relevant statistical structure into an iterative rather than sequential process. This is analogous to what has been done in the case of ‘clusterless decoding’ for spike sorting and decoding in neural spiking data with improved results [8]. In the case of EDA, this iterative process is successful because the underlying model is consistent with physiology. In addition, we compared our method to an established deconvolution-based EDA analysis paradigm and showed that our method is more consistent in pulse extraction and robust to noise. Finally, we have made the code open source, enabling others to use this methodology and further refine the technique. A preliminary version of this methodology was published in [9].

The balance of this paper is organized as follows. In Data and Methods, we outline how we used our insights about the physiology of sweat glands to design a robust pulse selection paradigm. Then in Results, we illustrate the use of this paradigm on two different subject cohorts and compare it to an established EDA analysis method. Finally, in the Discussion and Conclusion, we summarize our models and framework and the implications.

## II. Data and Methods

### A. Experimental data and preprocessing

For this study, we used EDA recordings from two experiments. The first is EDA we previously collected from 12 healthy volunteers between the ages of 22 and 34 while awake and at rest. The study was approved by the Massachusetts Institute of Technology (MIT) Institutional Review Board. Approximately one hour of EDA data was collected at 256 Hz from electrodes connected to the second and fourth digits of each subject’s non-dominant hand. Subjects were seated upright and instructed to remain awake. They were allowed to read, meditate, or watch something on a laptop or tablet, but not to write with the instrumented hand. One subject’s data were not included in the analysis because we learned after completing the experiment that the subject occasionally experienced a Raynaud’s type phenomenon, which would affect the quality of the EDA data. Data from the remaining 11 subjects were analyzed.

The second subject cohort consists of EDA recorded during a study of propofol-induced unconsciousness from eleven healthy volunteers while under propofol sedation [10]. The protocol was approved by the Massachusetts General Hospital (MGH) Human Research Committee. For all subjects, approximately 3 hours of data were recorded while using target-controlled infusion protocol. The data collection is described in detail in [10]. The infusion rate was increased and then decreased in a total of ten stages of roughly equal lengths to achieve target propofol concentrations of: 0 mg/kg/hr, 1, 2, 3, 4, 5, 3.75, 2.5, 1.25, 0. The two subject cohorts were collected in different years by different study teams and using different recording equipment systems. All data were analyzed using Matlab R2019a. All code will be available on the Neuroscience Statistics Research Laboratory website (www.neurostat.mit.edu).

Preprocessing consisted of two major steps, 1) detecting and removing artifacts and 2) isolating the phasic component. Both have been described in previously in [2]. Because of the level of high frequency noise seen in the recording equipment used for the propofol data, those data were additionally low-pass filtered with a cutoff of 3 Hz after artifact removal.

### B. Pulse extraction

We used a measure of locally-adjusted peak amplitude called prominence to account for the changing background filling level of the sweat glands. To compute prominence for each peak, we used the *findpeaks* algorithm in Matlab. This algorithm adjusts the amplitude of each peak in the signal as the height above the highest of neighboring “valleys” on either side. The valleys are chosen based on the lowest point in the signal between the peak and the next intersection with the signal of equal height on either side.

### C. Motivation for physiology-informed pulse selection

The goal of pulse selection from EDA data is to extract the set of pulses as close as possible to the ground truth of true EDA phasic activity. Therefore, relying on known properties of sweat gland physiology is key to distinguishing between pulses and noise. We previously showed that the bursting of sweat glands can be modeled as an integrate-and-fire process, which manifests as temporal structure that looks inverse Gaussian or like mixtures of inverse Gaussians in the inter-pulse intervals [2][3]. One way to characterize this is using tail behavior, which captures the heaviness of the tail of a right-skewed distribution using the ultimate settling rate. The ultimate settling rate is computed as the limit of the hazard function as *x* tends to infinity [11]. Different right-skewed distributions, such as the inverse Gaussian, lognormal, gamma, and exponential, differ in their tail behavior properties [12]. A slower settling rate indicates a heavier tail. Fitting a variety of models allows us to sample a range of tail behaviors (both lighter and heavier) which may represent mixtures of inverse Gaussians. Our previous work showed that EDA data favors heavier tailed models such as the lognormal and inverse Gaussian over lighter tailed models such as the gamma and exponential, likely due to the presence of regions of low phasic activity with long inter-pulse intervals [2]. Using these insights, we proceeded with the analysis in three phases for each subject cohort, detailed in the following sections. Overall, our approach aims to define a novel way of assessing to what degree pulses extracted from EDA data are consistent with known physiology.

### D. Phase I: Individual subject trends

We screened a range of prominence thresholds from 0.001 to 0.02 in increments of 0.001 and 0.02 to 0.08 in increments of 0.005 for each subject in the cohort. These limits were chosen to span from the smallest detectable deflection to the largest pulses in EDA data. At each prominence threshold, a set of pulses was extracted, from which four inter-pulse interval models were fit (the inverse Gaussian, lognormal, gamma, and exponential) by maximum likelihood [13]. For each, the Kolmogorov-Smirnov (KS) distance was computed as a measure of goodness-of-fit. The KS distance measures the maximum distance between the theoretical and empirical inter-pulse intervals after they have been rescaled using the time-rescaling theorem [14]. A smaller KS distance indicates a better fit. We also computed a 95% significance cutoff for the KS distance based on the number of pulses extracted [15]. A KS distance greater than this cutoff indicates that the data differ significantly from the model. This process resulted in four model KS-distances and one significance cut-off at each prominence threshold for each subject.

### E. Phase II: Subject cohort as a whole

We plotted the median KS-distance and significance cut-off per prominence threshold across subjects for the whole cohort. The goal of Phase II was to understand the effects of data collection settings, such as equipment used and experimental condition studied, on the EDA recordings from the subject cohort overall. We were specifically interested in the role of each of the four models to identify four specific trends which relate to the tail behavior of the four models:

1. The inverse Gaussian and/or lognormal models reach a “sweet spot” of being the best fitting distributions for some region of prominence thresholds. These models generally do not fit well at the lowest thresholds due to the presence of noise. The size of this “sweet spot” can vary from subject to subject but indicates that the pulse extraction is preserving the desired statistical structure without too much noise. This is the ideal region from which to select the optimal prominence threshold.
2. The exponential model is generally the worst fitting model for the majority of thresholds since it does not capture the physiology of sweat glands. However, with increasing threshold, the number of pulses decreases and therefore the significance cutoff becomes more generous. If the KS distance of the exponential model is above the significance cutoff at lower thresholds, the point at which it crosses under the significance cutoff marks the threshold at which there are now too few pulses to draw any conclusions. If the exponential model is under the significance cutoff even at lower thresholds (yet it is still the worst fitting model), this rule cannot be used to assess whether the number of pulses is sufficient.
3. The gamma model can be used to identify noise. The gamma model generally fits best in the presence of strong noise, usually at lower thresholds. This is because the presence of noise generally makes inter-pulse intervals short, which favors light-tailed inter-pulse interval distributions. The gamma is the model with the most flexibility in the tail and is the only one of the four models able to capture these very light tails. However, as noise is removed with increasing threshold, the KS-distance of the gamma model increases and “crosses” the lognormal and/or inverse Gaussian, as they fit the data better. We hereby refer to this trend (fitting well at lower thresholds and then slowly “rising out” with increasing threshold) as the “gamma signature”.
4. The gamma signature maybe be shifted left or right in a subject cohort, indicating the presence of lower or higher levels of noise overall, usually due to recording equipment or experimental conditions. For example, the gamma model may start with a relatively poor fit at lower thresholds, then improve to being the best fit model before slowly rising out. This is simply the gamma signature shifted to the right, which indicates that the subject cohort has higher levels of noise. The “sweet spot” for optimal thresholds will also be at higher thresholds for the subjects in the cohort.

We illustrate the methods with an analysis of a single subject for 3 different thresholds and the 4 candidate models (Fig. 2).

**Fig. 1.**
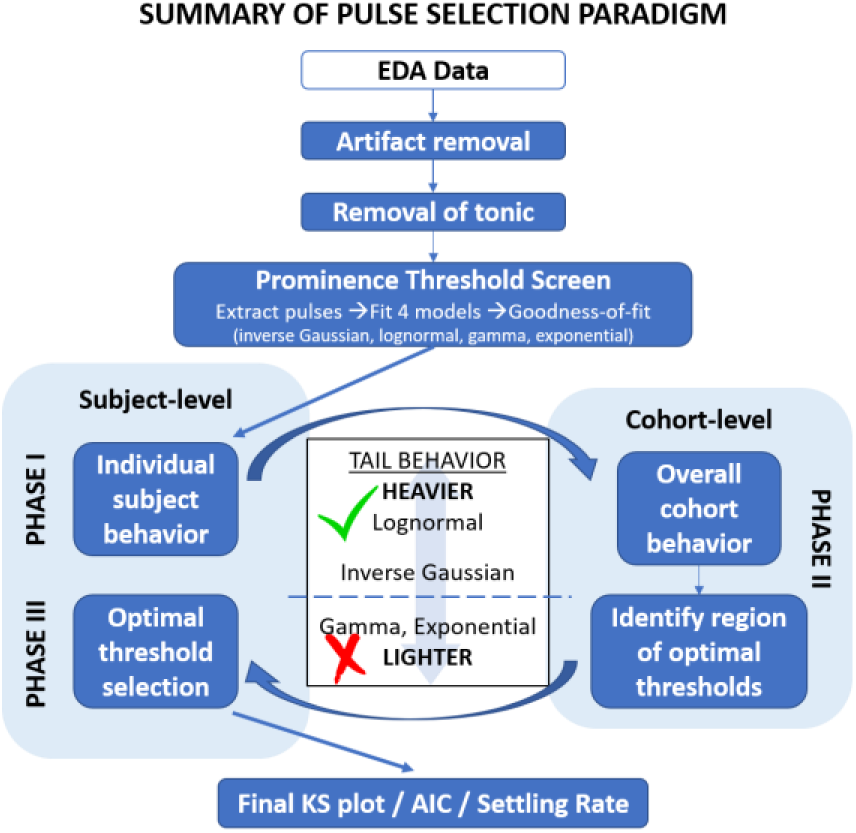
A schematic of the methods, starting from raw EDA data to verifying the goodness-of-fit of the chosen prominence threshold. In the center, the tail behavior properties of the 4 models are summarized.

**Fig. 2.**
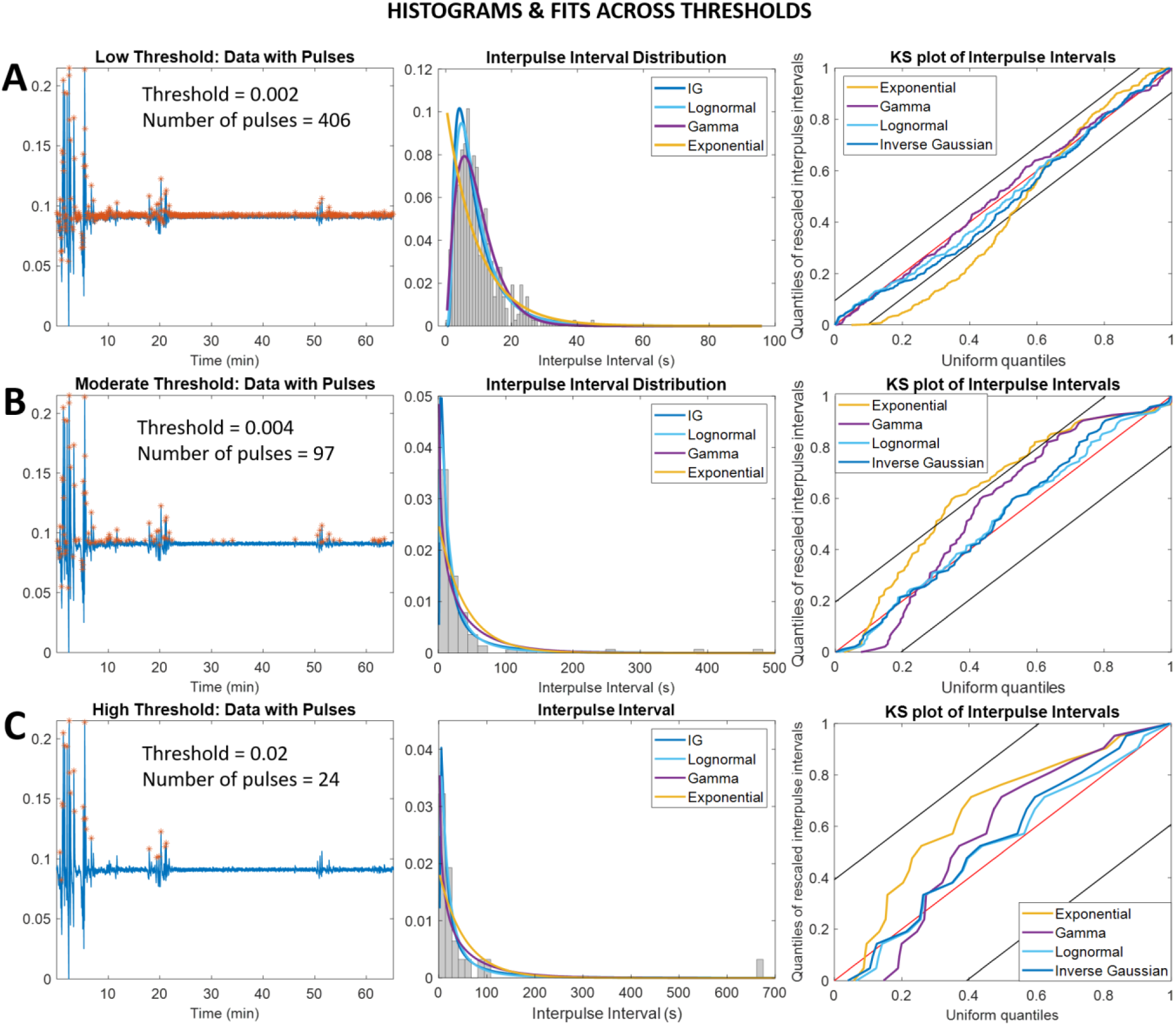
For a single subject’s data (S6), the threshold for pulse selection was gradually increased. At each of three representative thresholds, (a) 0.002 (low), (b) 0.004 (moderate), and (c) 0.02 (high), the pulses extracted, shape of the inter-pulse interval distribution, and the goodness-of-fit of the 4 distributions are shown.

At the lowest threshold (Fig. 2A), there are many short inter-pulse intervals due to the presence of noise in pulse selection, and therefore the inter-pulse interval distribution has almost no tail. The gamma model is best able to accommodate this. At moderate thresholds with less noise (Fig. 2B), the inverse Gaussian and lognormal are the best fit models since the inter-pulse interval distribution has a heavier tail. And finally, at the highest threshold (Fig. 2C), pulse extraction clearly excludes several pulses, and there are too few pulses to accurately characterize the inter-pulse interval distribution. The significance cutoffs are very generous, making all of the models appear to fit well. For each subject cohort, once each of these 4 trends had been identified, we proceeded to Phase II.

### F. Phase III: Optimal threshold selection per subject

The KS-distances and significance cut-offs across prominence thresholds computed in Phase I were plotted per subject. The trends were then compared to the overall cohort-level trends observed in Phase II. For a given subject, if all models behaved similarly across prominence thresholds in terms of goodness-of-fit with respect to the significance cutoff, the optimal prominence threshold was chosen from the “sweet spot” identified in Phase II, in which the inverse Gaussian and lognormal models are the best fitting.

If the models did not behave similarly to the cohort-level trends identified in Phase II, the subject was considered a potential outlier. In some cases, the same trends were seen, but shifted to the right (or left), which was an indication of higher (or lower) levels of noise in that subject’s data compared to the rest of the cohort in general. In those cases, the optimal threshold for that subject is likely to be much higher (or lower) than for the rest of the cohort. Finally, after selecting the optimal threshold for each subject, we made a full KS plot with 95% confidence bounds to verify that our pulse extraction and choice of threshold did indeed capture the statistical structure as intended. If the KS plot of a model follows the 45-degree diagonal and stays fully within the confidence bounds, then that model fits the data well. We computed the AIC and KS-distance for all models for each subject as well as the tail settling rate.

### G. Comparison to existing EDA algorithm

We extracted pulses from the awake and at rest cohort using an existing EDA analysis pipeline from the Leda Lab [16][17]. This algorithm is based on a deconvolution approach, in which EDA is assumed to be the result of the convolution of a neural input signal with a single impulse response function for each subject, which represents that subject’s sweat gland response to a single neural impulse. By recommendation of the authors, we used a continuous deconvolution analysis with the default parameters provided, which includes a peak threshold of 0.001 for all data. For each subject, after pulse extraction, we made a KS plot to assess whether the extracted pulses recovered statistical structure.

## III. Results

### A. Awake and At Rest Subject Cohort Phase I

The overall goodness-of-fit curves for the four models for the awake and at rest cohort (Fig. 3) show four trends previously mentioned. The lognormal becomes the best-fitting model around a threshold of 0.004, whereas the inverse Gaussian crosses the gamma around a threshold of 0.008. After this point, the lognormal and inverse Gaussians remain the best fitting models. The exponential starts out as and remains the worst-fitting model throughout, except for a very small region near a threshold of 0.017. The exponential crosses under the significance cutoff around a threshold of 0.01. The gamma distribution starts as the best-fitting model and crosses both the lognormal and inverse Gaussian. The gamma signature does not appear shifted to the right. Based on all four of these results, it seems that for the majority of subjects in this cohort, the optimal prominence threshold should be in the range of 0.004 to 0.01.

**Fig. 3.**
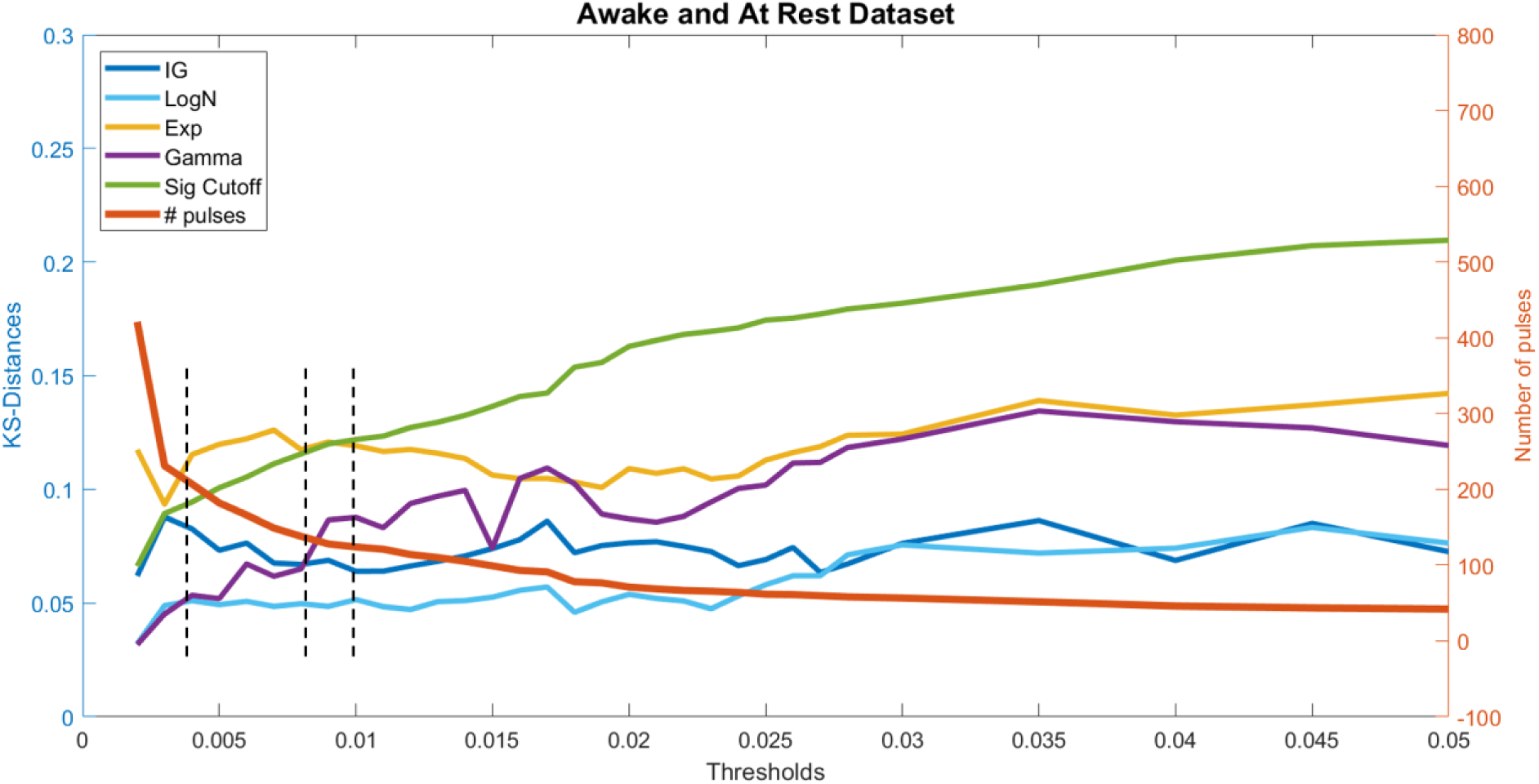
Overall goodness-of-fit curves for all 4 distributions (left y-axis) and number of pulses extracted (right y-axis) across the awake and at rest cohort, dashed vertical lines indicate, from left to right, (1) where the lognormal crosses under the gamma, (2) where the inverse Gaussian crosses under the gamma, and (3) where the exponential crosses under the significance cutoff

### B. Awake and At Rest Subject Cohort Phase II

Subject S6 shows similar goodness-of-fit curves to the overall awake and at rest cohort (Fig. 4B). The gamma starts as the best-fitting model and then crosses the lognormal and inverse Gaussian models. The exponential is always the worst-fitting model and crosses below the significance cut-off around a prominence threshold of 0.02. The gamma signature is not shifted to the right. We chose an optimal prominence threshold of 0.004. At that threshold, the pulses selected include all larger pulses as well as some smaller pulses in regions of sparse activity (Fig. 4C). The final KS-plot (Fig. 4D) shows that the lognormal and inverse Gaussian models remain close to the 45-degree line and within 95% confidence bounds throughout, but neither the exponential nor the gamma model does.

**Fig. 4.**
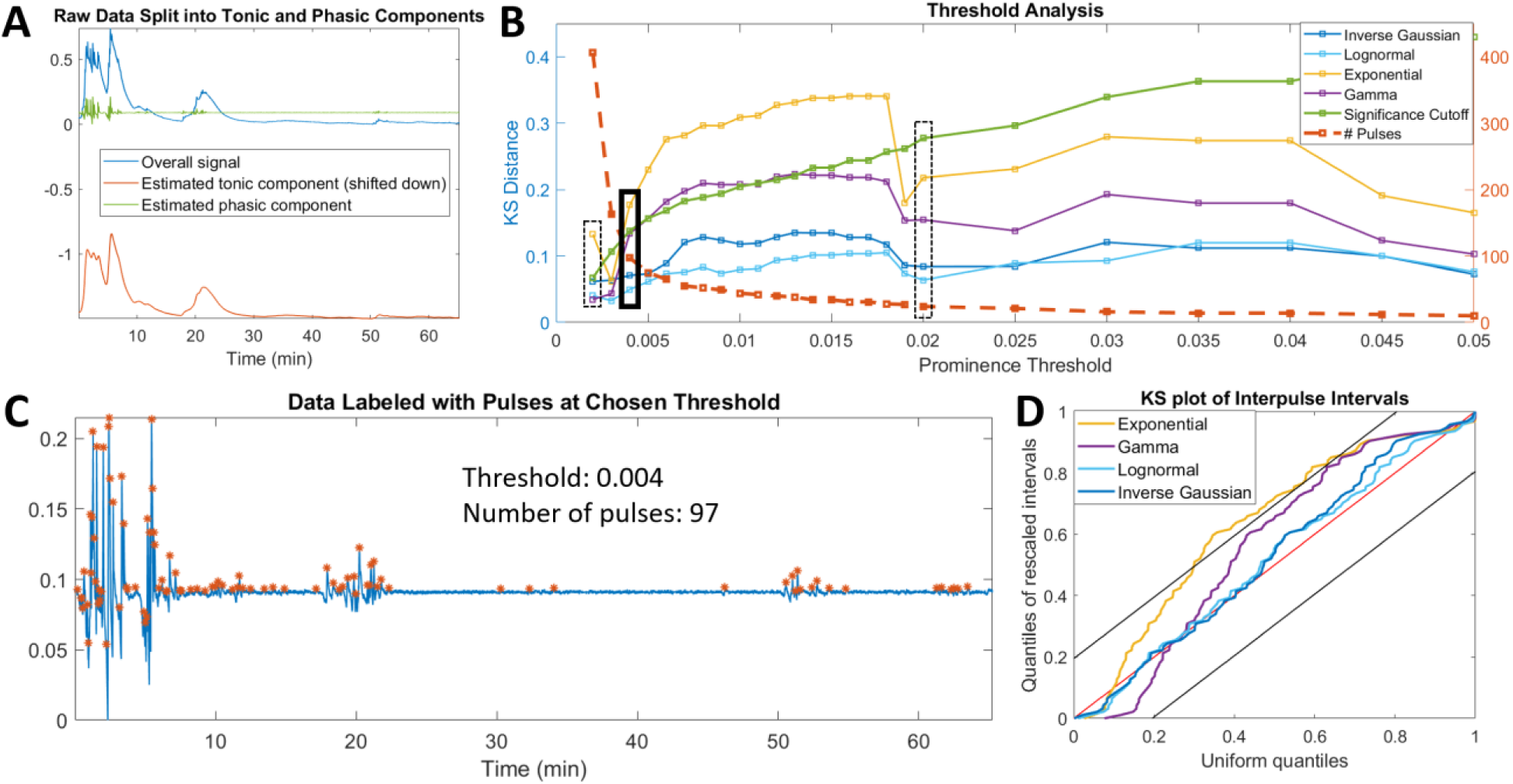
Results for Subject 6 from the awake and at rest cohort, showing agreement with the trends of the cohort as a whole. (a) Preprocessing of data by splitting into tonic and phasic components, (b) Screening of thresholds with chosen threshold marked with bolded rectangle, dashed rectangles indicate other thresholds (low and high) also shown in Fig. 1, (c) Pulses extracted at chosen threshold, (d) Full KS-plot showing goodness-of-fit at chosen threshold

Across the 11 subjects in the awake and at rest cohort, the final prominence thresholds used ranged between 0.0025 and 0.027. Nine of these 11 subjects had optimal thresholds between 0.004 and 0.01, as suggested from the full cohort analysis in Phase I. The total number of pulses in the one-hour time window ranged between 97 and 324, including the distantly spaced smaller pulses (Table I). The KS-distance and AIC results largely agreed with each other across subjects. At the respective optimal prominence thresholds for each subject, the lognormal or inverse Gaussian was the best fitting model for all 11 subjects according to KS-distance and 10 out of the 11 subjects according to AIC (Tables I and II). The exponential was the worst of the four models tested for 10 of the 11 subjects according to AIC and KS-distance. Specifically, with respect to KS-distance, the inverse Gaussian and lognormal models were within the significance cutoff for all subjects, the gamma for 8, but the exponential was only within the cutoff for 3 out of 11 subjects. These results suggest that our method was able to extract pulses while preserving the desired statistical structure.

**TABLE I.**
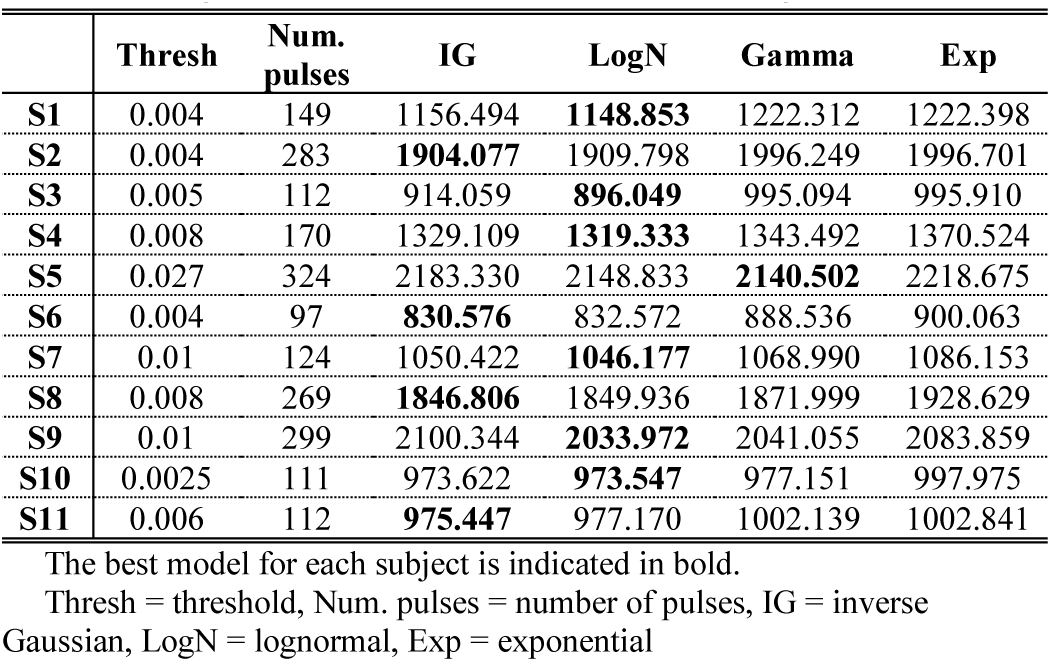
AIC Results for the Awake and At Rest Cohort

|  | Thresh | Num. pulses | IG | LogN | Gamma | Exp |
| --- | --- | --- | --- | --- | --- | --- |
| S1 | 0.004 | 149 | 1156.494 | **1148.853** | 1222.312 | 1222.398 |
| S2 | 0.004 | 283 | **1904.077** | 1909.798 | 1996.249 | 1996.701 |
| S3 | 0.005 | 112 | 914.059 | **896.049** | 995.094 | 995.910 |
| S4 | 0.008 | 170 | 1329.109 | **1319.333** | 1343.492 | 1370.524 |
| S5 | 0.027 | 324 | 2183.330 | 2148.833 | **2140.502** | 2218.675 |
| S6 | 0.004 | 97 | **830.576** | 832.572 | 888.536 | 900.063 |
| S7 | 0.01 | 124 | 1050.422 | **1046.177** | 1068.990 | 1086.153 |
| S8 | 0.008 | 269 | **1846.806** | 1849.936 | 1871.999 | 1928.629 |
| S9 | 0.01 | 299 | 2100.344 | **2033.972** | 2041.055 | 2083.859 |
| S10 | 0.0025 | 111 | 973.622 | **973.547** | 977.151 | 997.975 |
| S11 | 0.006 | 112 | **975.447** | 977.170 | 1002.139 | 1002.841 |

The best model for each subject is indicated in bold.
Thresh = threshold, Num. pulses = number of pulses, IG = inverse Gaussian, LogN = lognormal, Exp = exponential

**TABLE II.**
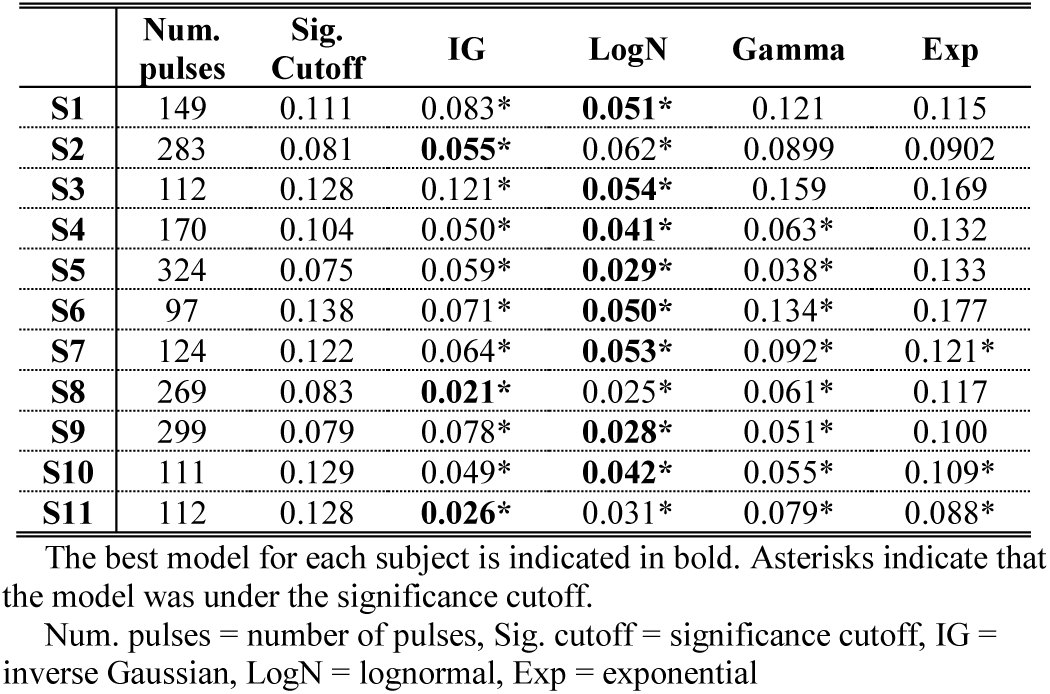
KS-Distance Results for the Awake and At Rest Cohort

Looking at the tail behavior analysis (Table III), the settling rate of the inverse Gaussian model is always less than that of the exponential and gamma, predicting a heavier tail than the exponential and gamma even though all three are commonly classified as medium-tailed distributions. The lognormal is commonly classified as a heavy-tailed distribution. Therefore, our method selects for a heavier tail in pulse extraction, representing heavy-tailed inverse Gaussians or mixtures of them which can be approximated by the lognormal.

**TABLE III.**
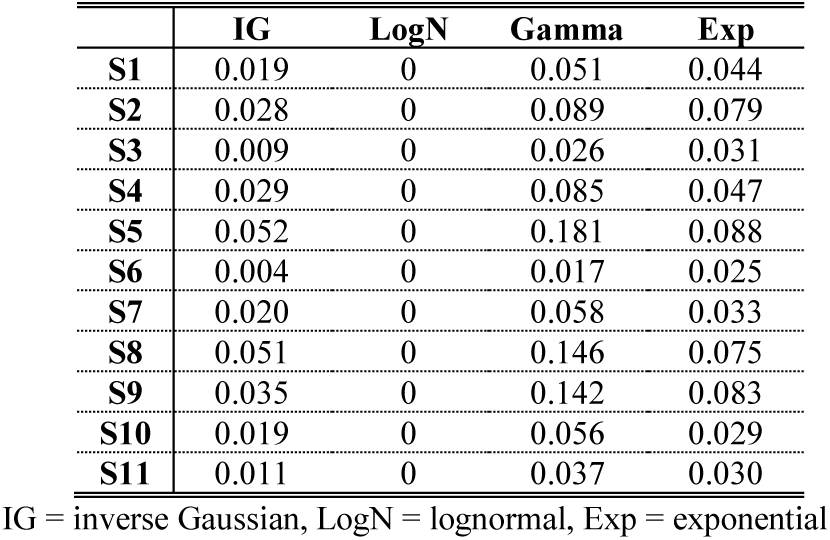
Settling Rate Results for the Awake and At Rest Cohort

Subject S5 presents an interesting case in which the goodness-of-fit curves deviate noticeably from the overall cohort trends (Fig. 5B). There are two clear deviations. First, the inverse Gaussian never crosses below the gamma model in goodness-of-fit, so the ‘sweet spot’ is determined by the lognormal alone. Second, the gamma signature appears shifted to the right, since it starts with a poor fit, then becomes the best fit model before crossing the lognormal. This also shifts the sweet spot and choice of optimal prominence threshold to the right. We accordingly chose an optimal threshold of 0.027. These trends would indicate that this subject’s data are characterized by a much higher level of noise than the rest of the cohort. This is also clear when looking at the data themselves (Fig. 5C). Interestingly, even though the KS distance selects for the lognormal over the gamma, the gamma is the best fit model according to AIC (Tables I and II). The tail behavior analysis indicates that the gamma has a much higher settling rate for this subject than any of the other models, indicating that it is able to accommodate the higher degree of noise and therefore shorter inter-pulse intervals with a very light tail.

**Fig. 5.**
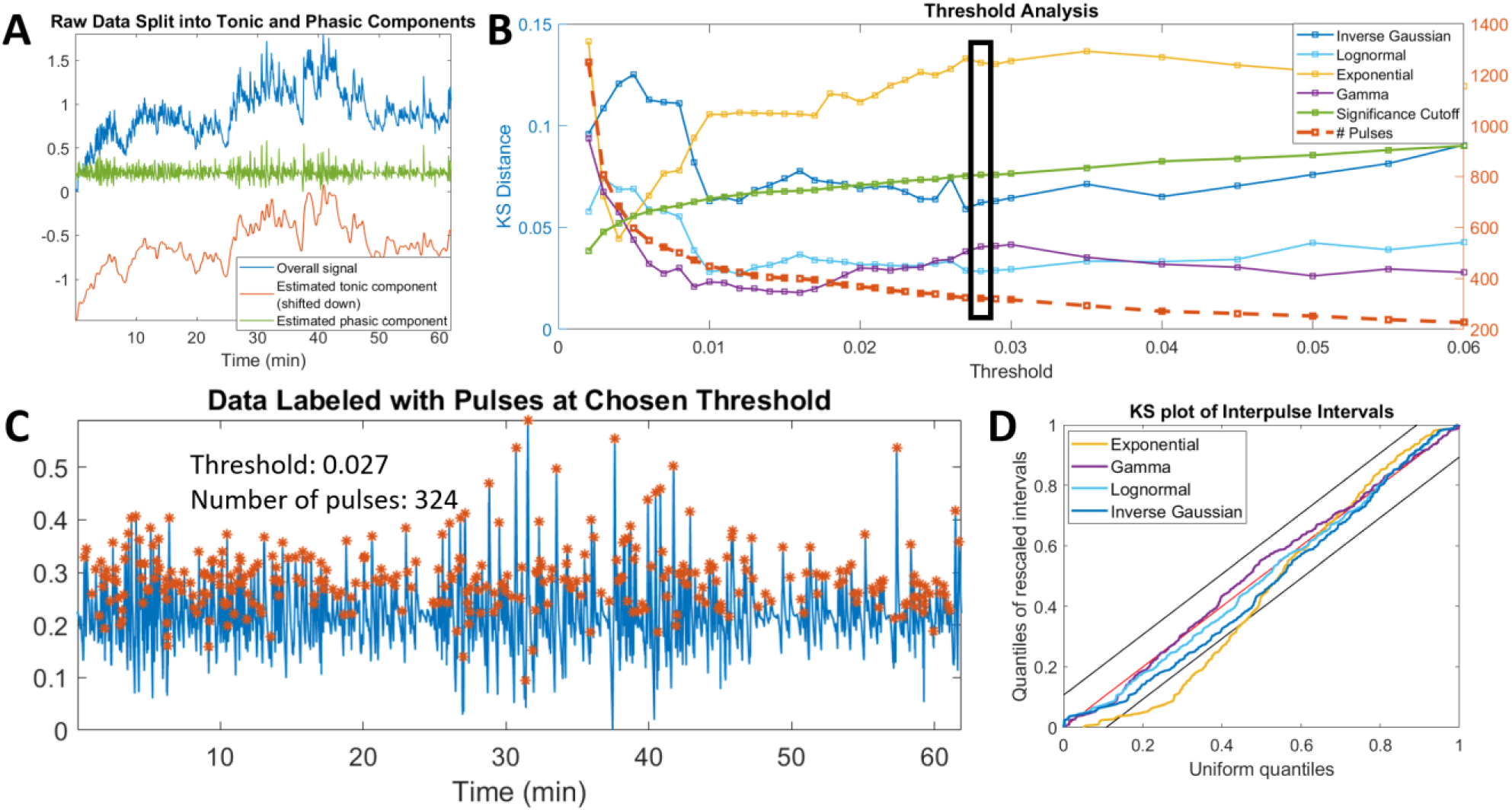
Results for Subject 5 from the awake and at rest cohort, showing noticeable differences from the trends of the cohort as a whole. (a) Preprocessing of data by splitting into tonic and phasic components, (b) Screening of thresholds with chosen threshold marked with bolded rectangle, (c) Pulses extracted at chosen threshold, (d) Full KS-plot showing goodness-of-fit at chosen threshold

### C. Propofol Sedation Subject Cohort Phase I

The overall goodness-of-fit curves for all four models for the propofol sedation cohort (Fig. 6) show some similarities to the awake and at rest cohort and some key differences with respect to the four trends previously discussed. Here, the lognormal and inverse Gaussian are always the best fitting models, largely remaining under the significance cut-off throughout. The exponential is almost always the worst-fitting model, but never crosses below the significance cutoff. The behavior of the gamma is the trend most distinctly different from that of the other cohort. The gamma is never the best-fitting model for this cohort nor is it ever under the significance cutoff. This reflects more stringent significance cutoffs and narrower 95% confidence bounds due to a longer duration of data (and therefore more data points) for each subject compared to the awake and at rest cohort. However, the shape of the gamma signature is still visible, just shifted to the right. The gamma starts with poor fit, then improves slightly to a local minimum around a threshold of 0.004 before slowly rising out. This suggests that the propofol sedation cohort has a much higher level of noise compared to the other, and therefore the region of optimal prominence thresholds will also be higher. Based on the Phase I analysis, the optimal prominence thresholds for most subjects in this cohort should be in the range of 0.015 to 0.055, where the gamma has risen out of its local minimum sufficiently and the exponential is still the worst fitting model. Because of the nature of the trends in this cohort, this range is very broad.

**Fig. 6.**
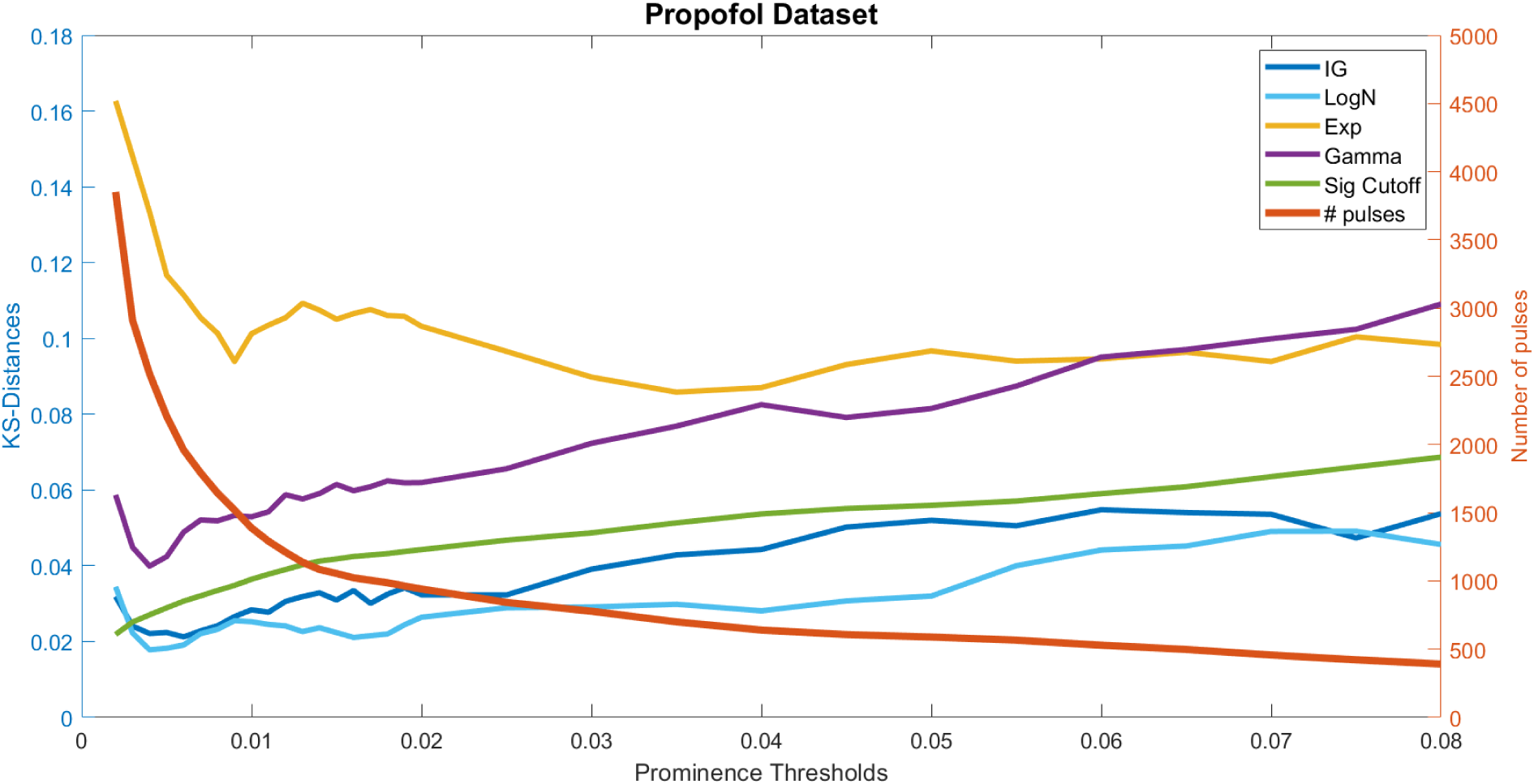
Overall goodness-of-fit curves for all 4 distributions (left y-axis) and number of pulses extracted (right y-axis) across the propofol sedation cohort

### D. Propofol Sedation Subject Cohort Phase II

Subject P09 shows similar goodness-of-fit curves to the overall propofol sedation cohort (Fig. 7B). The lognormal and inverse Gaussian models are the best fitting throughout. The gamma model signature is shifted to the right slightly, with it reaching a local minimum around a threshold of 0.01 and then rising out, even crossing above the exponential model, which is overall the worst fitting model throughout. We chose an optimal prominence threshold of 0.04. At that threshold, the extracted pulses include all pulse-like activity by visual inspection. Because of the level of noise inherent in the subject cohort overall, it is likely that the extracted pulses may include some noise. However, there are clear regions with no pulses that are still excluded correctly, such as between 20 and 60 minutes (Fig. 7C). In addition, the small zoom-in shows that the extracted pulses do actually correspond to pulse-like activity and not noise (Fig. 7C). The final KS-plot (Fig. 7D) shows that the lognormal and inverse Gaussian models stay close to the 45-degree line throughout while the gamma and exponential models do not.

**Fig. 7.**
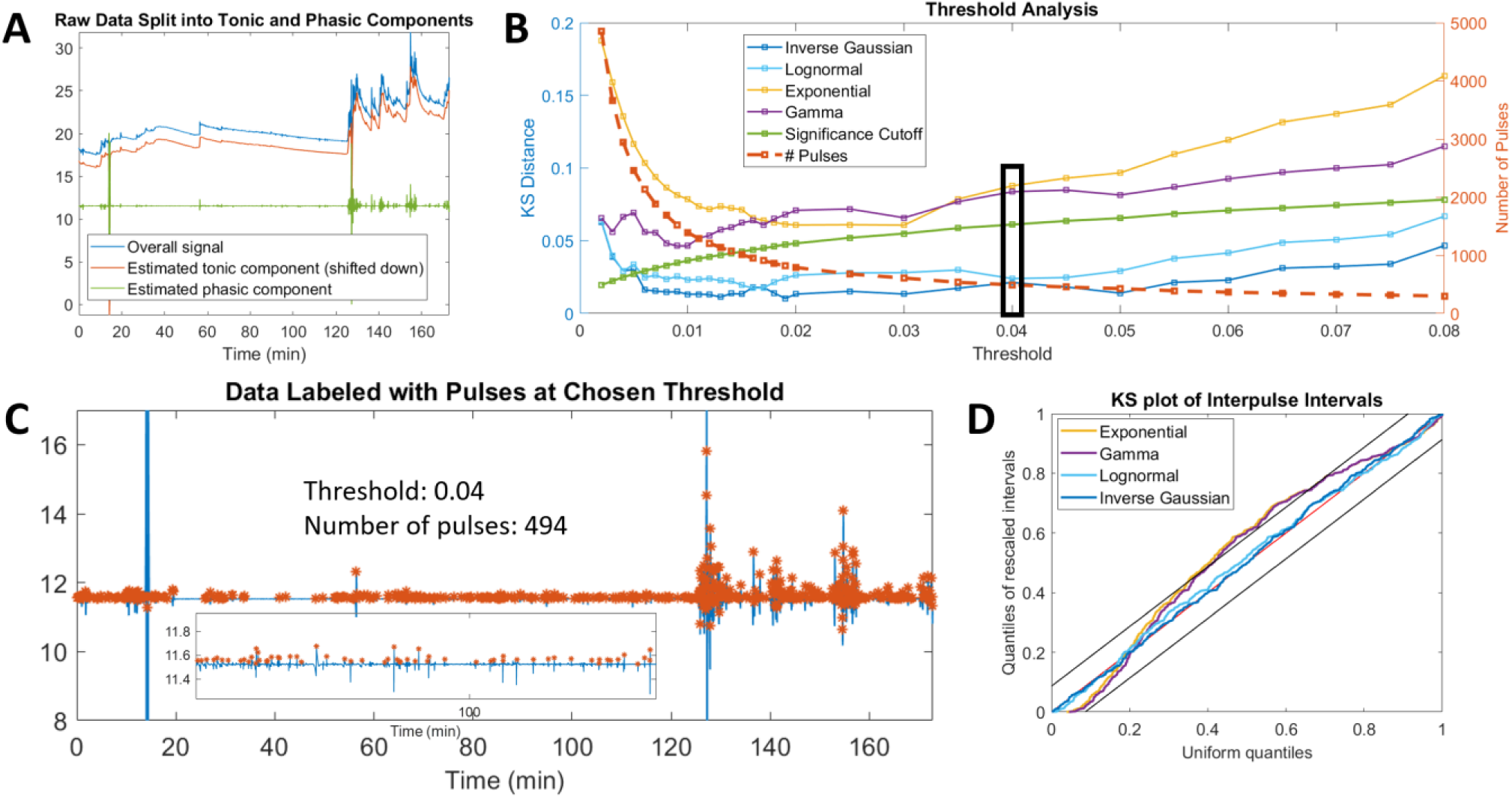
Results for Subject 8 from the propofol cohort, showing agreement with the trends of the cohort as a whole. (a) Preprocessing of data by splitting into tonic and phasic components, (b) Screening of thresholds with chosen threshold marked with bolded rectangle, (c) Pulses extracted at chosen threshold, (d) Full KS-plot showing goodness-of-fit at chosen threshold

Across the 11 subjects in the propofol sedation cohort, the final prominence thresholds used ranged between 0.02 and 0.055 (Table IV), which is fully within the range suggested from the cohort analysis in Phase I. The total number of pulses in the 3 to 4-hour time window ranged between 383 and 1250, which also reflects the higher degree of noise in the cohort. At the respective optimal prominence thresholds for each subject, either the lognormal or inverse Gaussian was the best fitting model for all 11 subjects according to both AIC and KS-distance (Tables IV and S-I). The exponential was the worst of the four models tested for 8 of the 11 subjects according to AIC and KS-distance, with the gamma performing slightly worse for the other 3 subjects. Specifically, with respect to KS-distance, the inverse Gaussian and lognormal models were within the significance cutoff for all subjects, the gamma for only one, the exponential for none of the subjects. Based on the tail behavior analysis (Table S-II), similar to the awake and at rest cohort, the settling rate of the inverse Gaussian model is always markedly less than that of the exponential and gamma. These results verify that our method was able to extract pulses while capturing the desired inverse Gaussian-like structure in the propofol sedation cohort, a second cohort with very different properties from the awake and at rest cohort.

**TABLE IV.**
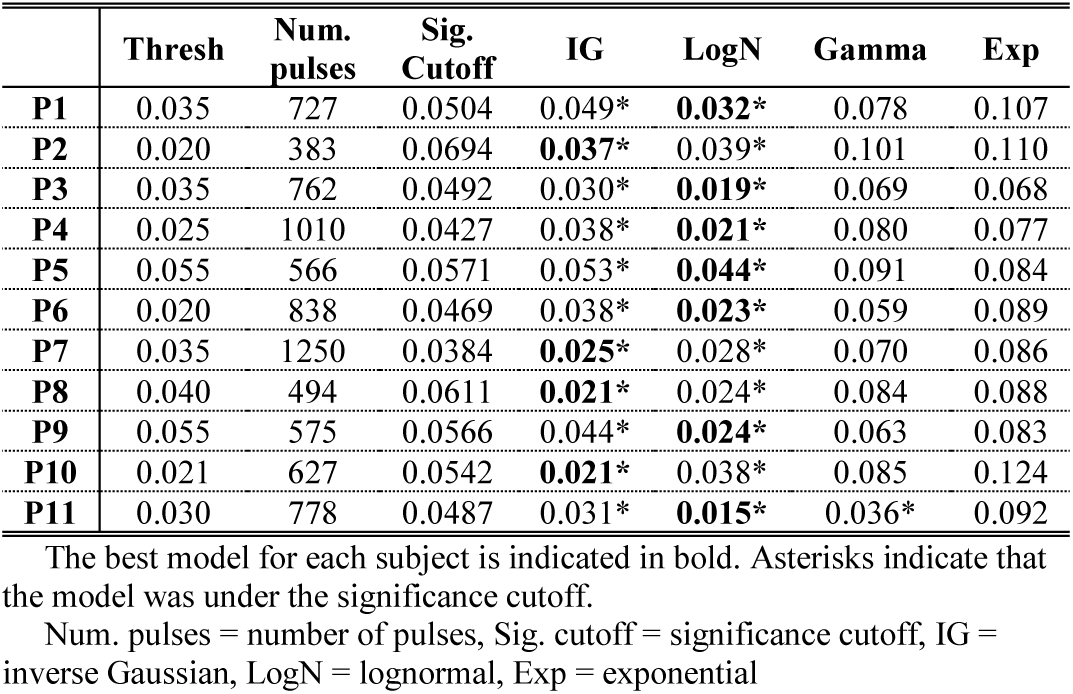
KS-Distance Results for the Propofol Sedation Cohort

### E. Comparison to the Ledalab algorithm

The Ledalab algorithm extracts almost an order of magnitude more pulses than our method for each subject in the awake and at rest cohort (Table S-III), likely due to the low default threshold value of 0.001 for pulses. In comparison, with our method, we chose a threshold at least 2.5 times greater for all subjects. Since this method does not adapt any parameters to the properties of the data at hand, there is no guarantee that pulse extraction will be consistent across subjects or cohorts. This is reflected in the statistical properties of the pulses extracted. The extracted pulses largely look like noise (Fig. 8A), and the goodness-of-fit analysis (Fig. 8B) indicates that for the majority of subjects, none of the four models offers an accurate statistical description of the data.

**Fig. 8.**
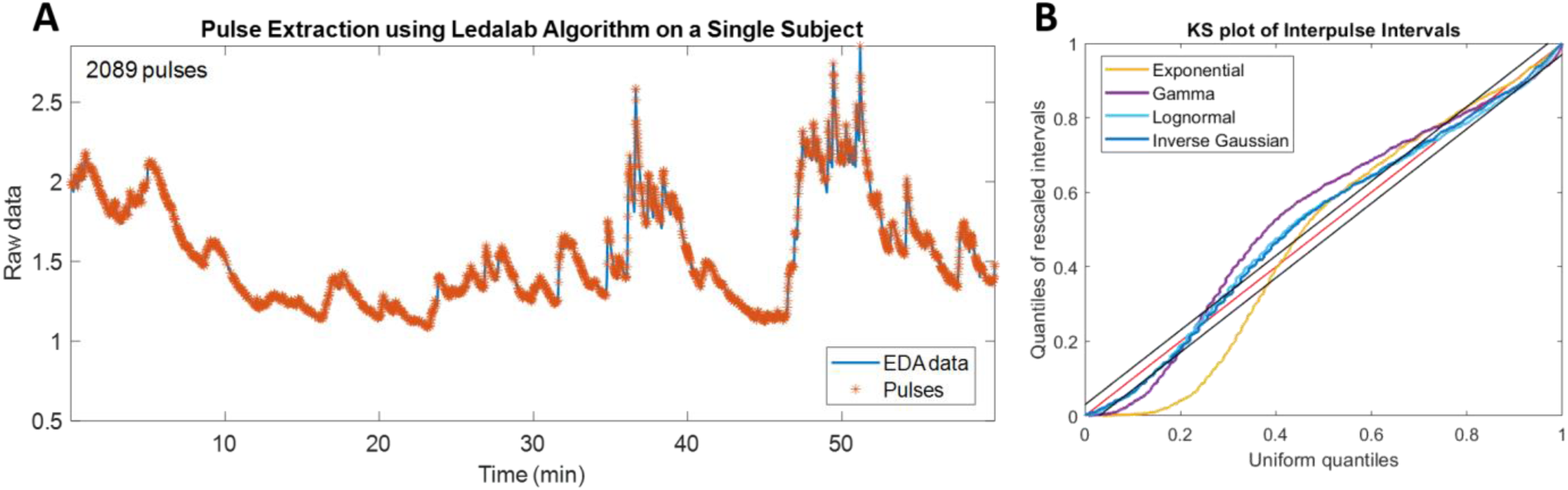
(a) Pulse selection and (b) goodness-of-fit results using the Ledalab algorithm on S8 from the awake and at rest cohort.

## IV. Discussion

In this work, we define a systematic and robust method to extract pulses from EDA data that preserves the statistical structure in the data derived from physiology while excluding noise. This method also allows for an assessment of the signal to noise profile across a whole cohort of data and identification of individual subjects whose data do not behave like the rest of the cohort. We tested this method on two cohorts, each collected using different equipment and under different experimental conditions. We showed that the method captures the statistical structure in both. In each cohort, the goodness-of-fit properties inherently reflect the signal to noise ratio in that cohort. Finally, we compared our method to pulse selection using the Ledalab algorithm for EDA and showed that unlike our method, it does not separate signal from noise in a new subject cohort on which it was not built.

Our framework for modeling and analyzing EDA data provides a structured and computationally efficient way to extract the relevant information from the data. We use a total of seven parameters across four models (inverse Gaussian, lognormal, gamma, and exponential) to determine how to extract pulses. By doing so, we reduce the arbitrariness of assigning a threshold to the data simply based on visual inspection or assigning a single threshold across all datasets. We also allow the structure in the data to inform pre-processing and analysis, instead of passing the data through an analysis pipeline and assessing the information the extracted data at the end.

The continuous deconvolution-based Ledalab algorithm, on the other hand, assumes a single unchanging pulse amplitude threshold across all subjects and cohorts. This results in too generous of pulse selection, with an average of one pulse every two seconds across all subjects (mean of 1880 pulses in one hour). This is far too frequent given known sweat gland physiology and numerous other studies of EDA. In addition, the goodness-of-fit results indicate that none of the models is under the significance cut-off for the majority of subjects at this threshold. This is similar to the threshold screening results of our method at very low thresholds in high-noise data. Even if one were to attempt to threshold the Ledalab pulse selection with a higher threshold, it must be done arbitrarily based on visual inspection alone. Our method uses the idea of an informed screen across thresholds without imposing assumptions about noise, pulse amplitude, or pulse shape across the cohort.

By developing an analysis pipeline that is computationally efficient, statistically rigorous, and physiology-based, we enable robust capture of relevant information from EDA data. We can also track the goodness-of-fit of our models in any setting. This robustness is a significant step forward in allowing EDA to be a clinical marker for sympathetic activity in diverse conditions such as pain, anxiety, depression, and sleep. Our future work will investigate models for capturing the relevant information in the amplitudes of the pulses as well and applying both types of information to understand dynamics. Code is available on our website (http://www.neurostat.mit.edu/). We hope to further refine this methodology and release further versions of the code that incorporate pulse amplitude analysis and other modifications based on the feedback we receive from those who use it on their own data.

## V. Conclusion

We are presenting a novel paradigm for analyzing EDA data to extract pulses based on the statistical characteristics of the EDA time series. Our paradigm performs consistently across different subject cohorts and verifies that the selected pulses reflect known sweat gland physiology, a process unique to our paradigm. Accurate pulse selection from EDA is key to inferring underlying sympathetic information. Sympathetic activation is implicated in many physiologic and psychological states, including stress, anxiety, depression, sleep, and pain. Our physiology-based paradigm for robust pulse selection from EDA data opens the door for EDA to be part of the clinical standard for measurement and tracking in these states.

## Acknowledgment

S. S. would like to thank the staff of the Massachusetts Institute of Technology Clinical Research Center.

## Notes

This work was supported by funds from the Picower Institute for Learning and Memory, the National Science Foundation Graduate Research Fellowship Program, and NIH Award P01-GM118629 (to E.N.B.).

### Competing Interest Statement

The authors have declared no competing interest.

### Summary of Updates

Minor updates to the text, figures, and figure captions throughout. Supplemental files updated.

## References

[1] W. Boucsein, Electrodermal Activity. New York, NY: Springer, 2012.

[2] S. Subramanian, R. Barbieri, E.N. Brown. (2020). Point process temporal structure characterizes electrodermal activity, bioRxiv (2020).

[3] S. Subramanian, R. Barbieri, E.N. Brown, “A point process characterization of electrodermal activity,” in Proceedings of the 40th IEEE International Conference on Engineering in Medicine and Biology, Honolulu, HI, 2018.

[4] E. Schrodinger. (1915). Zur theorie der fall-und steigversuche an teilchen mit brownscher bewegung. Physikalische Zeitschrift. 16.

[5] G. Gerstein, B. Mandelbrot. (1964). Random walk models for the spike activity of a single neuron. Biophys. J. 4, 41–68.

[6] R. Chhikara, J. Folks, The Inverse Gaussian Distribution: Theory, Methodology, and Applications. New York, NY: Marcel Dekker, 1989.

[7] R. Barbieri, E. Matten, A. Alabi, E. Brown. (2005). A point-process model of human heartbeat intervals: new definitions of heart rate and heart rate variability. Am. J. Physiol. Hear. Circ. Physiol. 288, H424–435.

[8] X. Deng, D.F. Liu, K. Kay, L.M. Frank, U.T. Eden. (2015). Clusterless Decoding of Position from Multiunit Activity Using a Marked Point Process Filter, Neural computation. 27(7), 1438–60.

[9] S. Subramanian, R. Barbieri, E.N. Brown, “A systematic method for preprocessing and analyzing electrodermal activity,” in Proceedings of the 41st IEEE International Conference on Engineering in Medicine and Biology, Berlin, Germany, 2019.

[10] P.L. Purdon et al. (2013). Electroencephalogram signatures of loss and recovery of consciousness from propofol, PNAS 110, E1142–1151.

[11] L. Halliwell. (2013). Classifying the tails of loss distributions. Casualty Actuar. Soc. 2.

[12] J. Folks, R. Chhikara. (1978). The inverse gaussian distribution and its statistical application – a review. J. Royal Stat. Soc. Ser. B (Methodological). 40, 263–289.

[13] Y. Pawitan, In All Likelihood. Oxford, UK: Clarendon Press, 2013.

[14] E. Brown, R. Barbieri, V. Ventura, R. Kass, L. Frank. (2001). The time-rescaling theorem and its application to neural spike train data analysis. Neural Comput. 14, 325–346.

[15] D. Daley, D. Vere-Jones, An Introduction to the Theory of Point Processes: Volume II: General Theory and Structure. New York, NY: Springer, 2007.

[16] M. Benedek, C. Kaernbach. (2010). Decomposition of skin conductance data by means of nonnegative deconvolution. Psychophysiology 47, 647–658.

[17] M. Benedek, C. Kaernbach. (2010). A continuous measure of phasic electrodermal activity. J. Neurosci. Methods. 190, 80–91.

[18] H. Storm, et al. (2002). Skin conductance correlates with perioperative stress. Acta Anaesthesiol. Scand. 46, 887–895.

[19] H. Storm, M. Shafiei, K. Myre, J. Raeder. (2005). Palmar skin conductance compared to a developed stress score and to noxious and awakening stimuli on patients in anaesthesia. Acta Anaesthesiol. Scand. 49, 798–803.

[20] A. Sano, R. Picard, R. Stickgold. (2014). Quantitative analysis of wrist electrodermal activity during sleep. Int. J. Psychophysiol. 94, 382–389.

[21] R Faghih, et al., “Characterization of fear conditioning and fear extinction by analysis of electrodermal activity,” in Proceedings of the 37th IEEE International Conference on Engineering in Medicine and Biology, Milan, Italy, 2015.

[22] D. Alexander, et al. (2005). Separating individual skin conductance responses in a short interstimulus-interval paradigm. J. Neurosci. Methods 146, 116–123.

[23] A. Greco, G. Valenza, A. Lanata, E. Scilingo, L. Citi. (2016). A convex optimization approach to electrodermal activity processing. IEEE Transactions on Biomed. Eng. 63, 797–804.

