## Supplementary Material for "A Model-Based Approach for Pulse Selection from Electrodermal Activity"

#### AWAKE AND AT REST COHORT

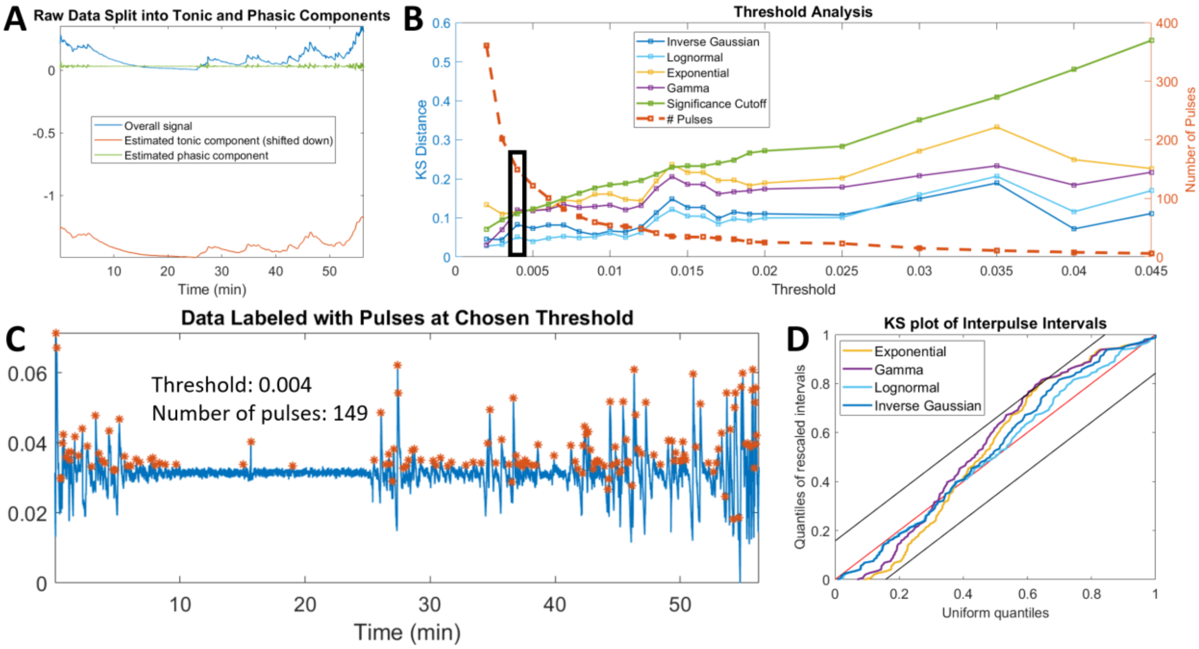


Fig. S1. Results for Subject 1 from the awake and at rest cohort, showing agreement with the trends of the cohort as a whole. (a) Preprocessing of data by splitting into tonic and phasic components, (b) Screening of thresholds with chosen threshold marked with bolded rectangle, (c) Pulses extracted at chosen threshold, (d) Full KS-plot showing goodness-of-fit at chosen threshold

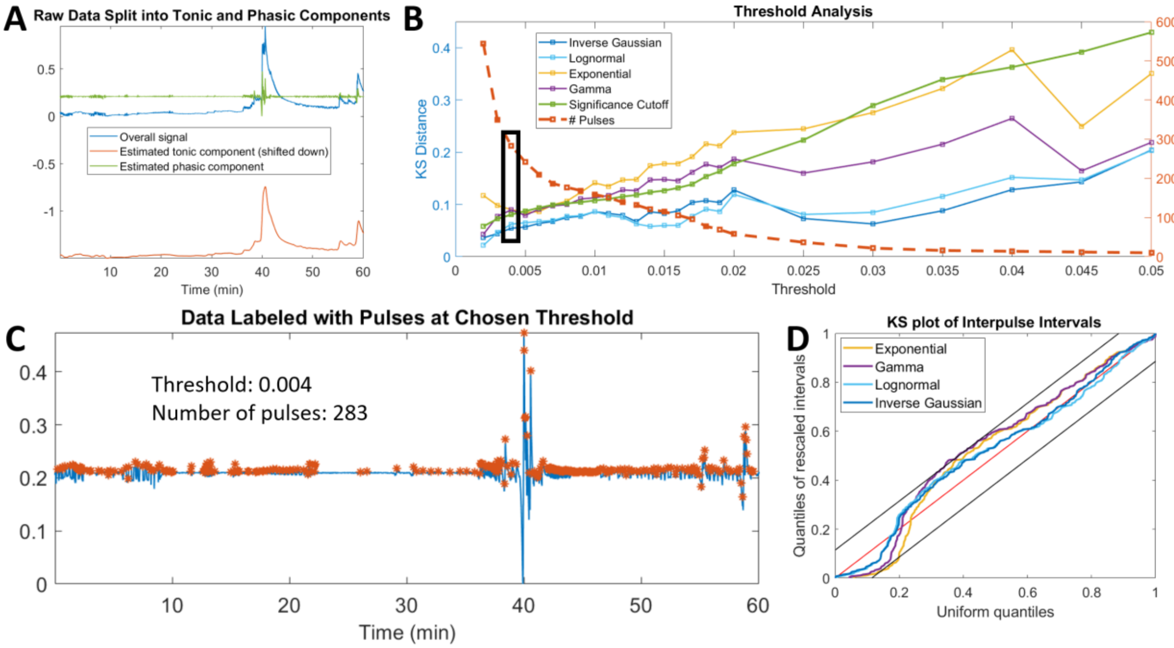


Fig. S2. Results for Subject 2 from the awake and at rest cohort, showing agreement with the trends of the cohort as a whole. (a) Preprocessing of data by splitting into tonic and phasic components, (b) Screening of thresholds with chosen threshold marked with bolded rectangle, (c) Pulses extracted at chosen threshold, (d) Full KS-plot showing goodness-of-fit at chosen threshold

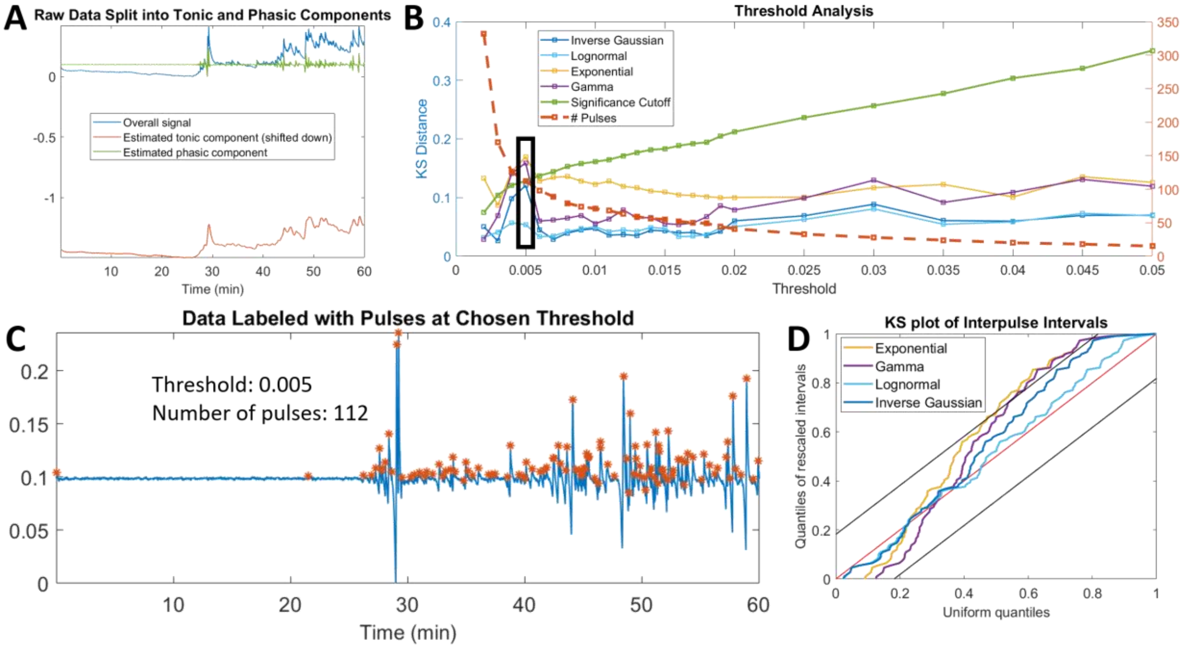


Fig. S3. Results for Subject 3 from the awake and at rest cohort, showing agreement with the trends of the cohort as a whole. (a) Preprocessing of data by splitting into tonic and phasic components, (b) Screening of thresholds with chosen threshold marked with bolded rectangle, (c) Pulses extracted at chosen threshold, (d) Full KS-plot showing goodness-of-fit at chosen threshold

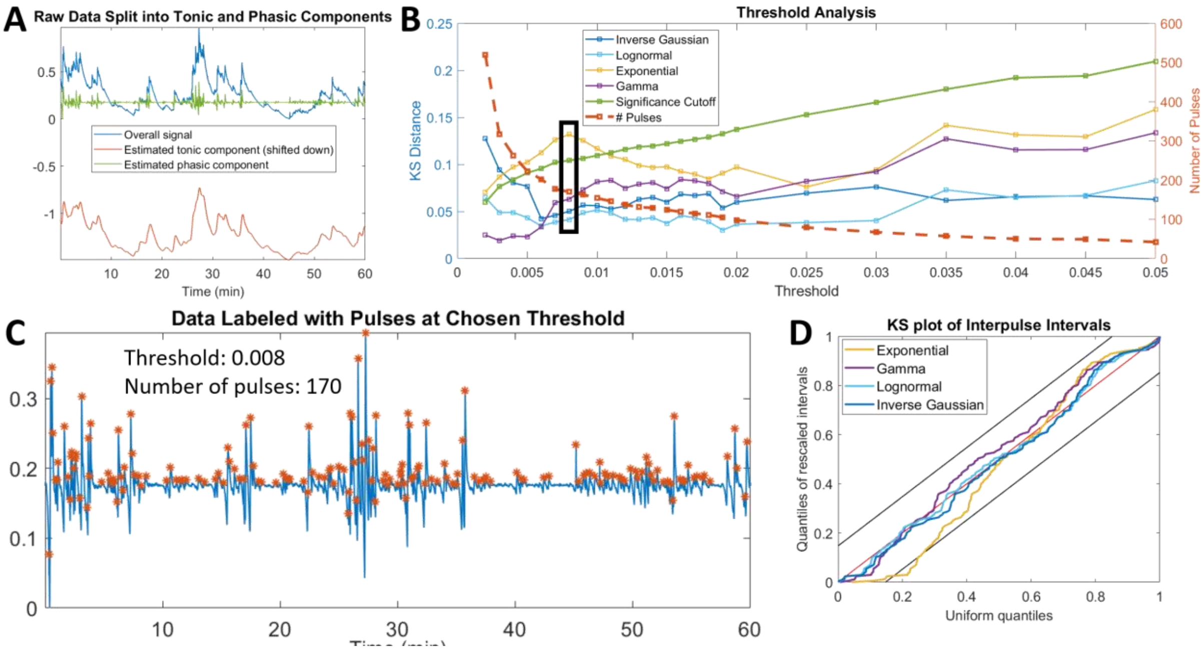


Fig. S4. Results for Subject 4 from the awake and at rest cohort, showing agreement with the trends of the cohort as a whole. (a) Preprocessing of data by splitting into tonic and phasic components, (b) Screening of thresholds with chosen threshold marked with bolded rectangle, (c) Pulses extracted at chosen threshold, (d) Full KS-plot showing goodness-of-fit at chosen threshold

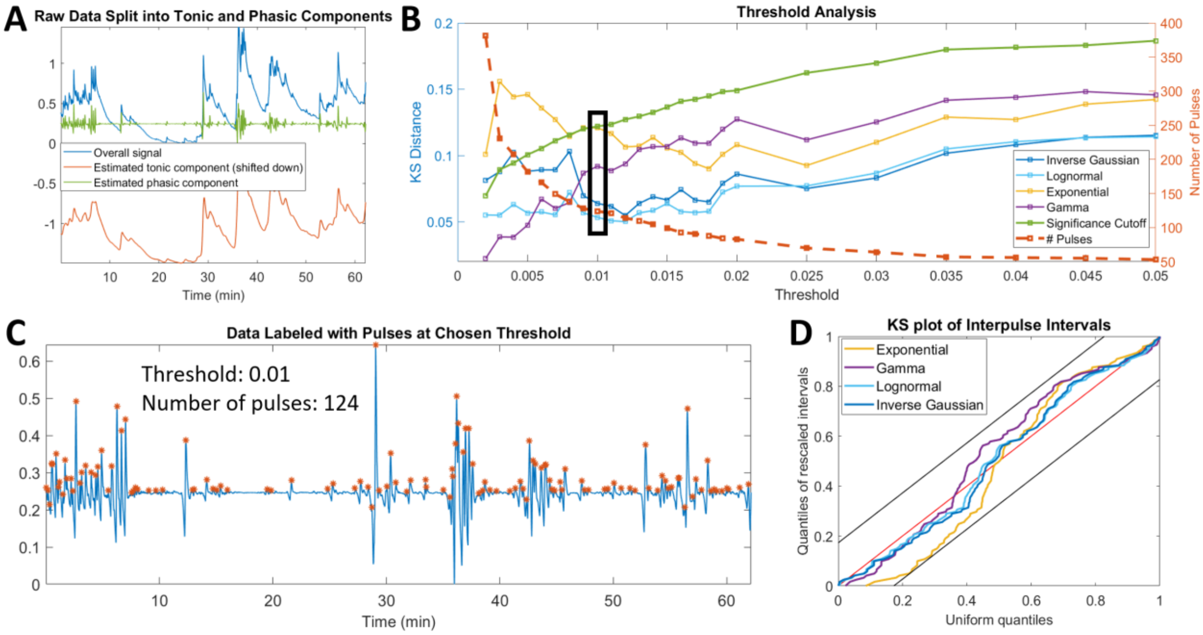


Fig. S5. Results for Subject 7 from the awake and at rest cohort, showing agreement with the trends of the cohort as a whole. (a) Preprocessing of data by splitting into tonic and phasic components, (b) Screening of thresholds with chosen threshold marked with bolded rectangle, (c) Pulses extracted at chosen threshold, (d) Full KS-plot showing goodness-of-fit at chosen threshold

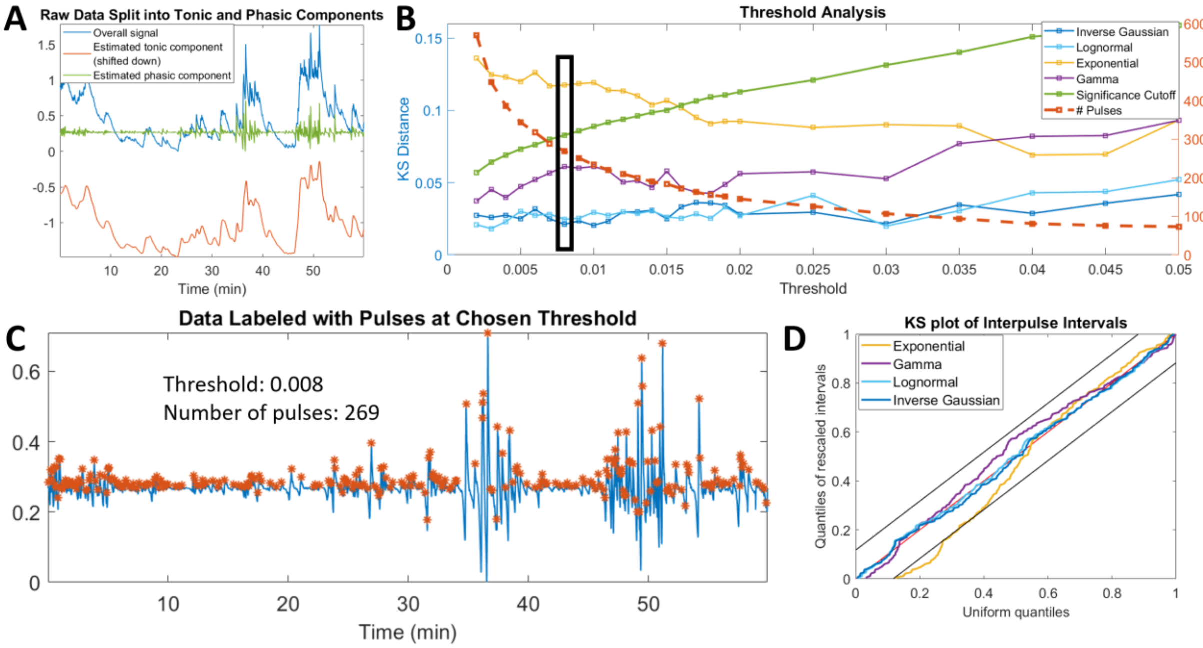


Fig. S6. Results for Subject 8 from the awake and at rest cohort, showing agreement with the trends of the cohort as a whole. (a) Preprocessing of data by splitting into tonic and phasic components, (b) Screening of thresholds with chosen threshold marked with bolded rectangle, (c) Pulses extracted at chosen threshold, (d) Full KS-plot showing goodness-of-fit at chosen threshold

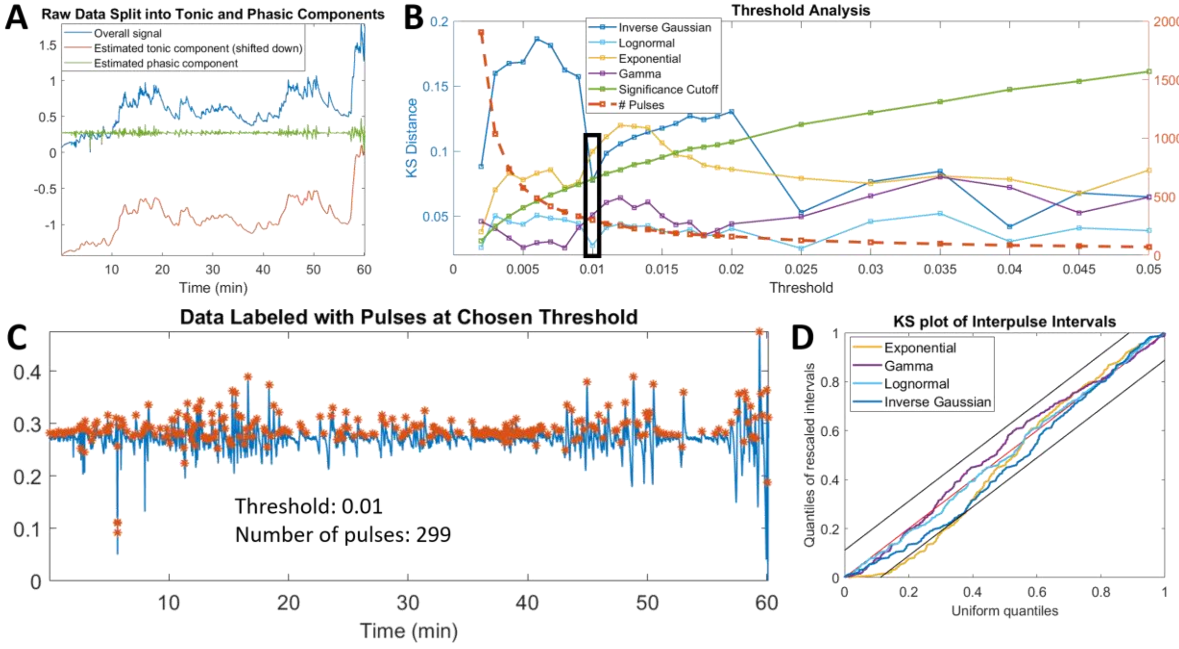


Fig. S7. Results for Subject 9 from the awake and at rest cohort, showing agreement with the trends of the cohort as a whole. (a) Preprocessing of data by splitting into tonic and phasic components, (b) Screening of thresholds with chosen threshold marked with bolded rectangle, (c) Pulses extracted at chosen threshold, (d) Full KS-plot showing goodness-of-fit at chosen threshold

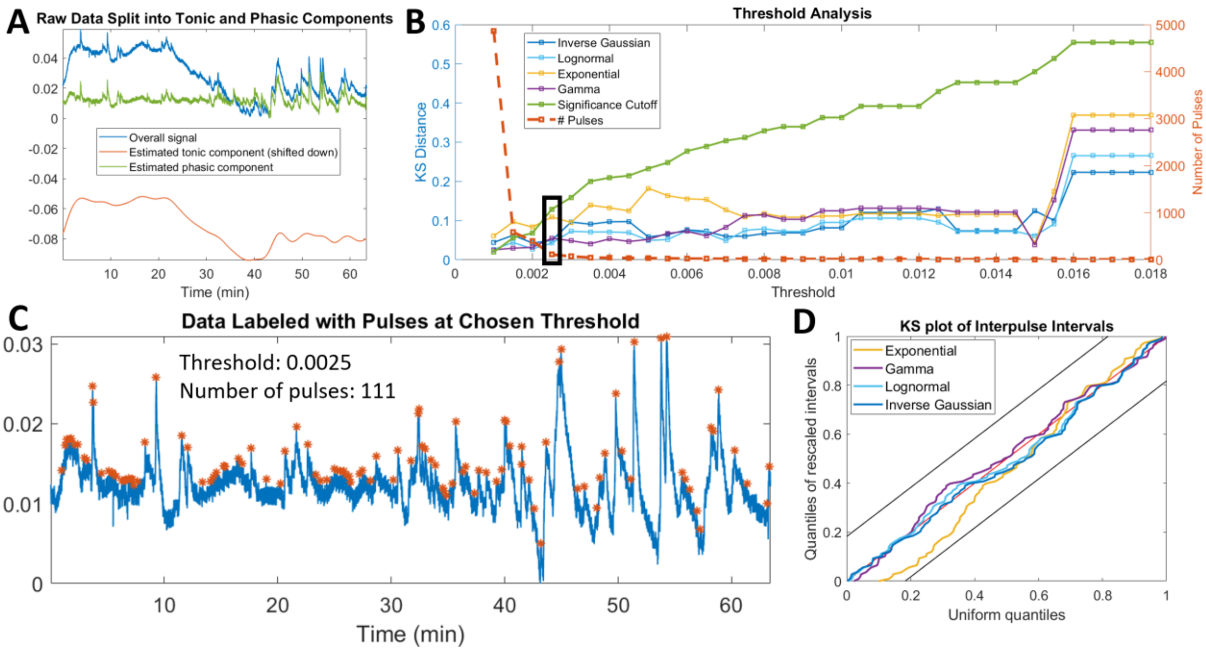


Fig. S8. Results for Subject 10 from the awake and at rest cohort, showing agreement with the trends of the cohort as a whole. (a) Preprocessing of data by splitting into tonic and phasic components, (b) Screening of thresholds with chosen threshold marked with bolded rectangle, (c) Pulses extracted at chosen threshold, (d) Full KS-plot showing goodness-of-fit at chosen threshold

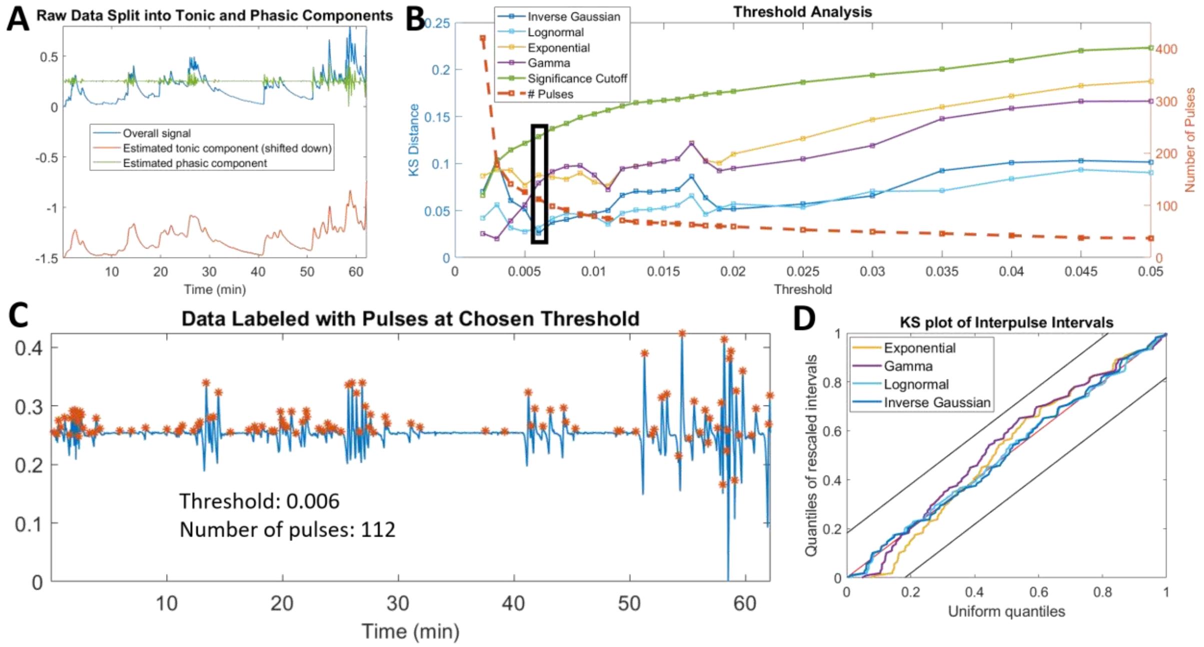


Fig. S9. Results for Subject 11 from the awake and at rest cohort, showing agreement with the trends of the cohort as a whole. (a) Preprocessing of data by splitting into tonic and phasic components, (b) Screening of thresholds with chosen threshold marked with bolded rectangle, (c) Pulses extracted at chosen threshold, (d) Full KS-plot showing goodness-of-fit at chosen threshold

### PROPOFOL SEDATION COHORT

TABLE S-I

AIC RESULTS FOR THE PROPOFOL SEDATION COHORT

| | Thresh | Num. pulses | IG | LogN | Gamma | Exp |
|---|---|---|---|---|---|---|
| **P1** | 0.035 | 727 | 4892.359 | **4862.670** | 5077.407 | 5159.771 |
| **P2** | 0.020 | 383 | **3232.026** | 3243.971 | 3370.773 | 3370.680 |
| **P3** | 0.035 | 762 | 5259.942 | **5259.534** | 5470.766 | 5468.826 |
| **P4** | 0.025 | 1010 | 7169.775 | **7130.224** | 7440.207 | 7440.261 |
| **P5** | 0.055 | 566 | 4293.377 | **4260.999** | 4400.896 | 4428.529 |
| **P6** | 0.020 | 838 | 5512.970 | **5490.560** | 5701.877 | 5772.100 |
| **P7** | 0.035 | 1250 | **7489.674** | 7498.297 | 7771.781 | 7866.955 |
| **P8** | 0.040 | 494 | **3837.628** | 3847.869 | 3991.711 | 3990.792 |
| **P9** | 0.055 | 575 | 4479.243 | **4448.334** | 4522.479 | 4570.867 |
| **P10** | 0.021 | 627 | **4617.829** | 4651.130 | 4874.572 | 4917.459 |
| **P11** | 0.030 | 778 | 5355.024 | **5332.838** | 5392.146 | 5523.467 |

The best model for each subject is indicated in bold.

Thresh = threshold, IG = inverse Gaussian, LogN = lognormal, Exp = exponential

TABLE S-II

SETTLING RATE RESULTS FOR THE PROPOFOL SEDATION COHORT

| | IG | LogN | Gamma | Exp |
|---|---|---|---|---|
| **P1** | 0.047 | 0 | 0.125 | 0.078 |
| **P2** | 0.008 | 0 | 0.030 | 0.033 |
| **P3** | 0.021 | 0 | 0.076 | 0.075 |
| **P4** | 0.020 | 0 | 0.072 | 0.068 |
| **P5** | 0.023 | 0 | 0.073 | 0.054 |
| **P6** | 0.045 | 0 | 0.129 | 0.087 |
| **P7** | 0.058 | 0 | 0.170 | 0.117 |
| **P8** | 0.012 | 0 | 0.045 | 0.048 |
| **P9** | 0.023 | 0 | 0.076 | 0.051 |
| **P10** | 0.009 | 0 | 0.039 | 0.054 |
| **P11** | 0.045 | 0 | 0.140 | 0.078 |

IG = inverse Gaussian, LogN = lognormal, Exp = exponential

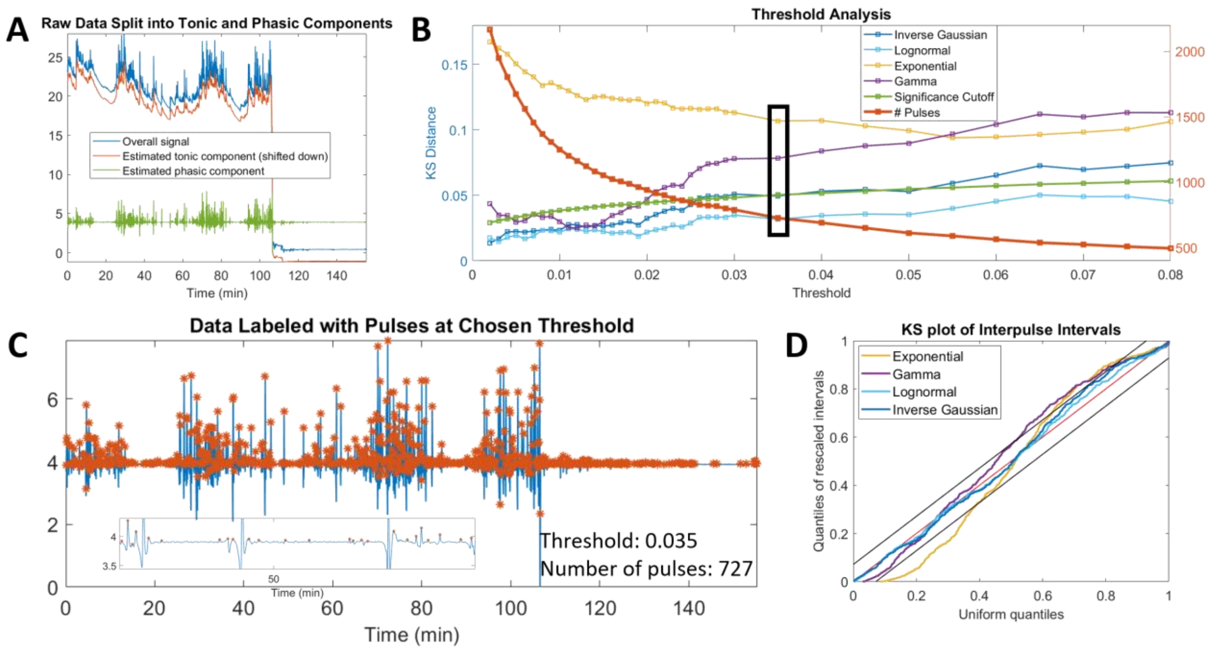


Fig. S10. Results for Subject 1 from the propofol sedation cohort, showing agreement with the trends of the cohort as a whole. (a) Preprocessing of data by splitting into tonic and phasic components, (b) Screening of thresholds with chosen threshold marked with bolded rectangle, (c) Pulses extracted at chosen threshold, (d) Full KS-plot showing goodness-of-fit at chosen threshold

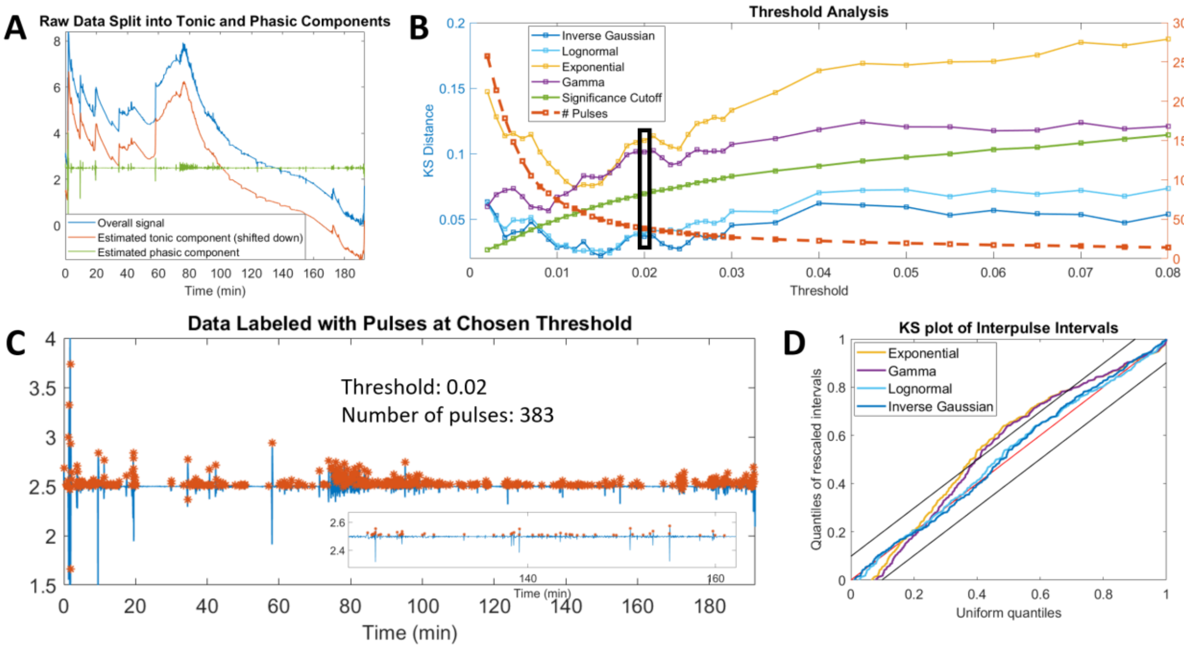


Fig. S11. Results for Subject 2 from the propofol sedation cohort, showing agreement with the trends of the cohort as a whole. (a) Preprocessing of data by splitting into tonic and phasic components, (b) Screening of thresholds with chosen threshold marked with bolded rectangle, (c) Pulses extracted at chosen threshold, (d) Full KS-plot showing goodness-of-fit at chosen threshold

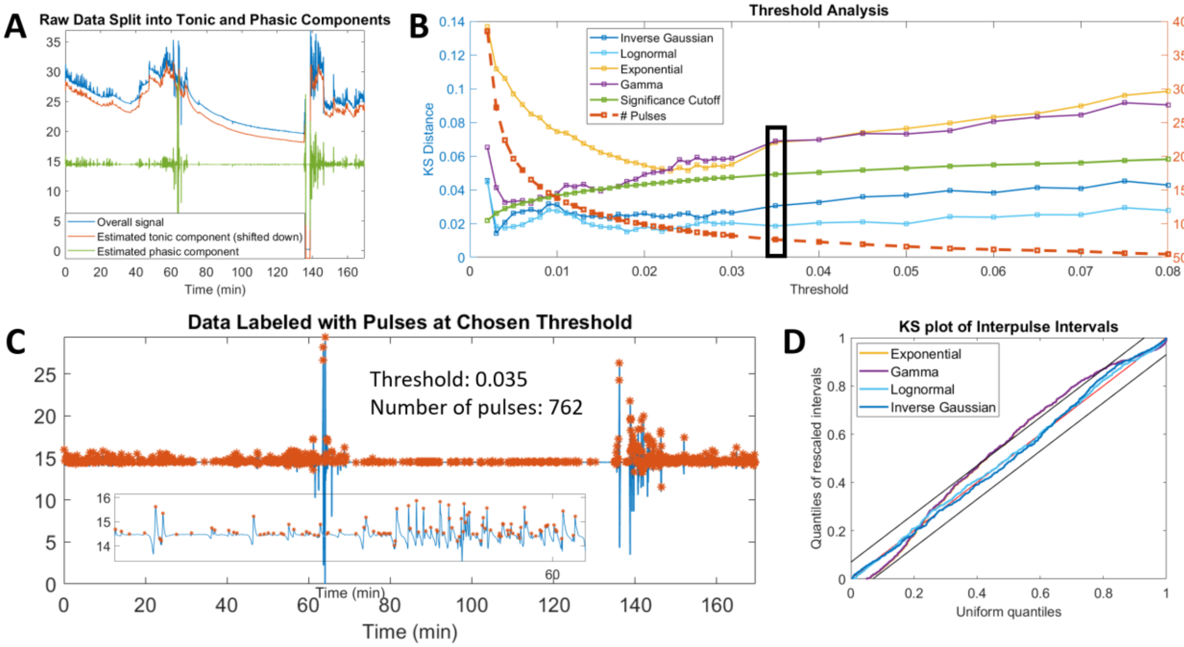


Fig. S12. Results for Subject 3 from the propofol sedation cohort, showing agreement with the trends of the cohort as a whole. (a) Preprocessing of data by splitting into tonic and phasic components, (b) Screening of thresholds with chosen threshold marked with bolded rectangle, (c) Pulses extracted at chosen threshold, (d) Full KS-plot showing goodness-of-fit at chosen threshold

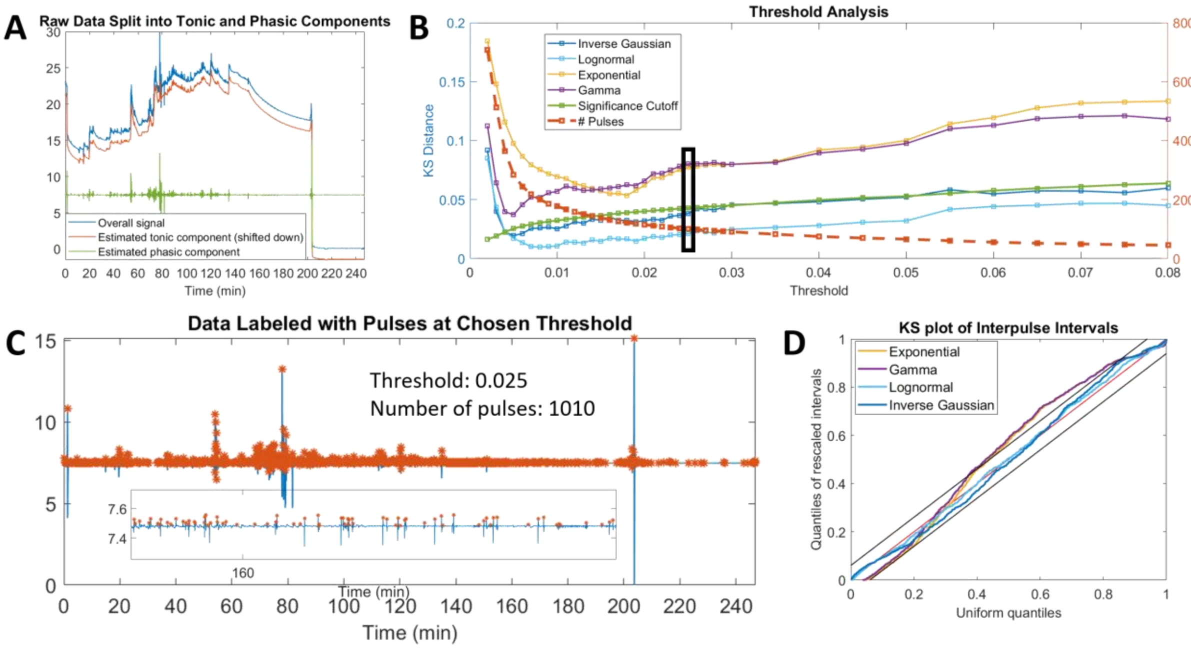


Fig. S13. Results for Subject 4 from the propofol sedation cohort, showing agreement with the trends of the cohort as a whole. (a) Preprocessing of data by splitting into tonic and phasic components, (b) Screening of thresholds with chosen threshold marked with bolded rectangle, (c) Pulses extracted at chosen threshold, (d) Full KS-plot showing goodness-of-fit at chosen threshold

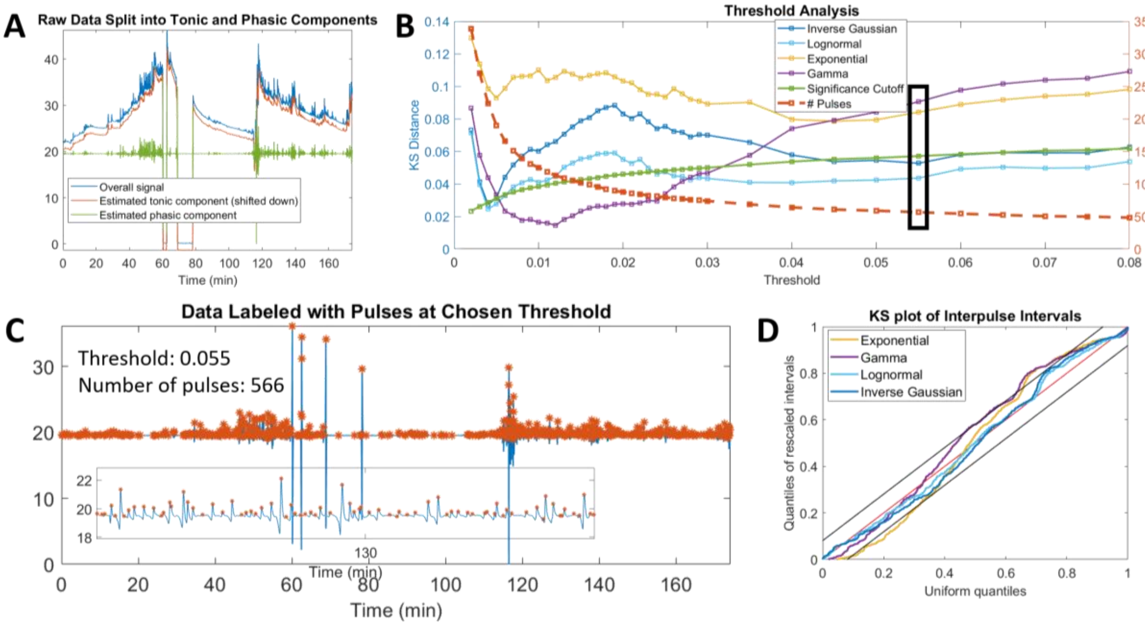


Fig. S14. Results for Subject 5 from the propofol sedation cohort, showing agreement with the trends of the cohort as a whole. (a) Preprocessing of data by splitting into tonic and phasic components, (b) Screening of thresholds with chosen threshold marked with bolded rectangle, (c) Pulses extracted at chosen threshold, (d) Full KS-plot showing goodness-of-fit at chosen threshold

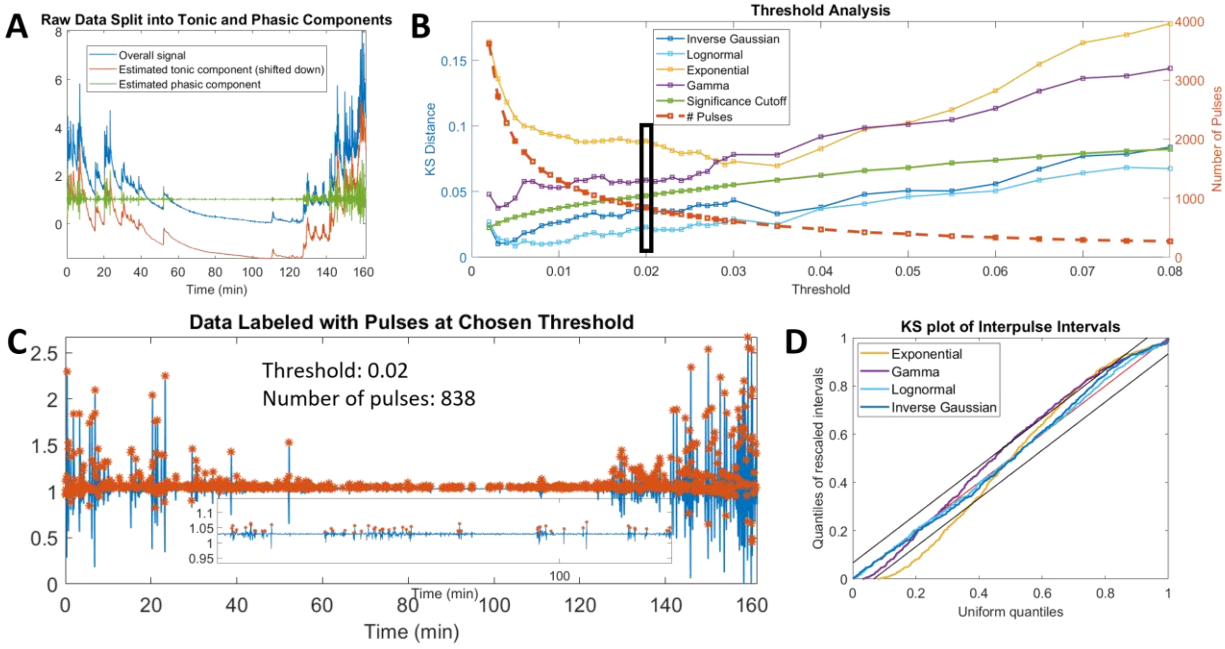


Fig. S15. Results for Subject 6 from the propofol sedation cohort, showing agreement with the trends of the cohort as a whole. (a) Preprocessing of data by splitting into tonic and phasic components, (b) Screening of thresholds with chosen threshold marked with bolded rectangle, (c) Pulses extracted at chosen threshold, (d) Full KS-plot showing goodness-of-fit at chosen threshold

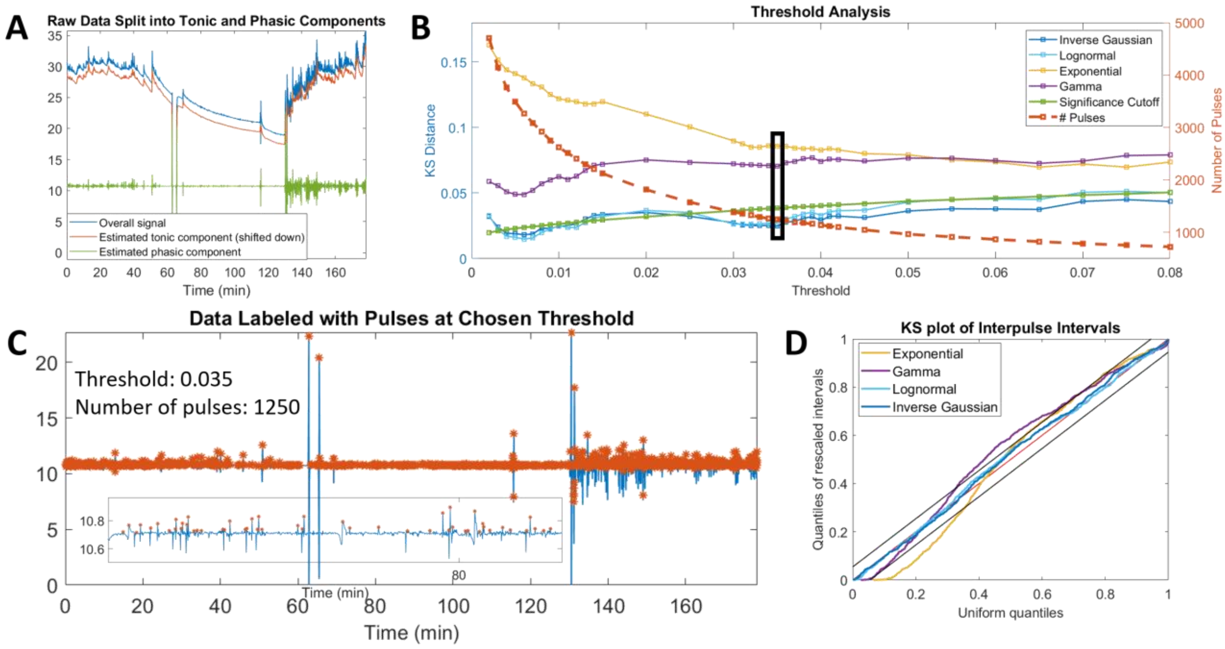


Fig. S16. Results for Subject 7 from the propofol sedation cohort, showing agreement with the trends of the cohort as a whole. (a) Preprocessing of data by splitting into tonic and phasic components, (b) Screening of thresholds with chosen threshold marked with bolded rectangle, (c) Pulses extracted at chosen threshold, (d) Full KS-plot showing goodness-of-fit at chosen threshold

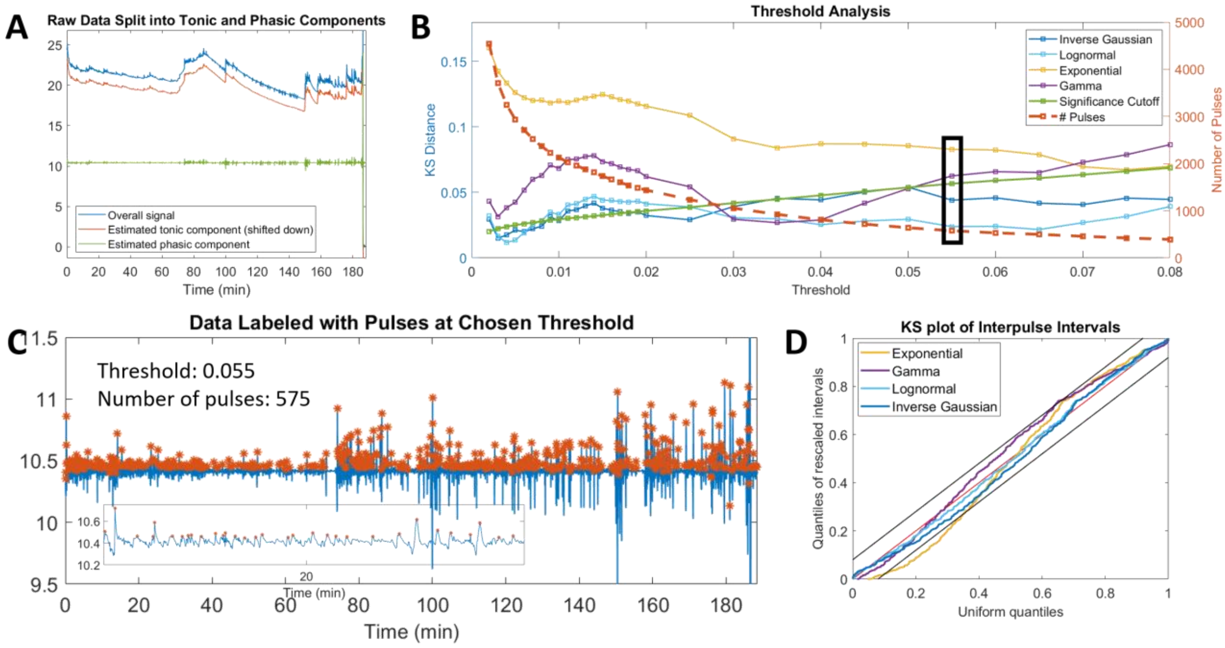


Fig. S17. Results for Subject 9 from the propofol sedation cohort, showing agreement with the trends of the cohort as a whole. (a) Preprocessing of data by splitting into tonic and phasic components, (b) Screening of thresholds with chosen threshold marked with bolded rectangle, (c) Pulses extracted at chosen threshold, (d) Full KS-plot showing goodness-of-fit at chosen threshold

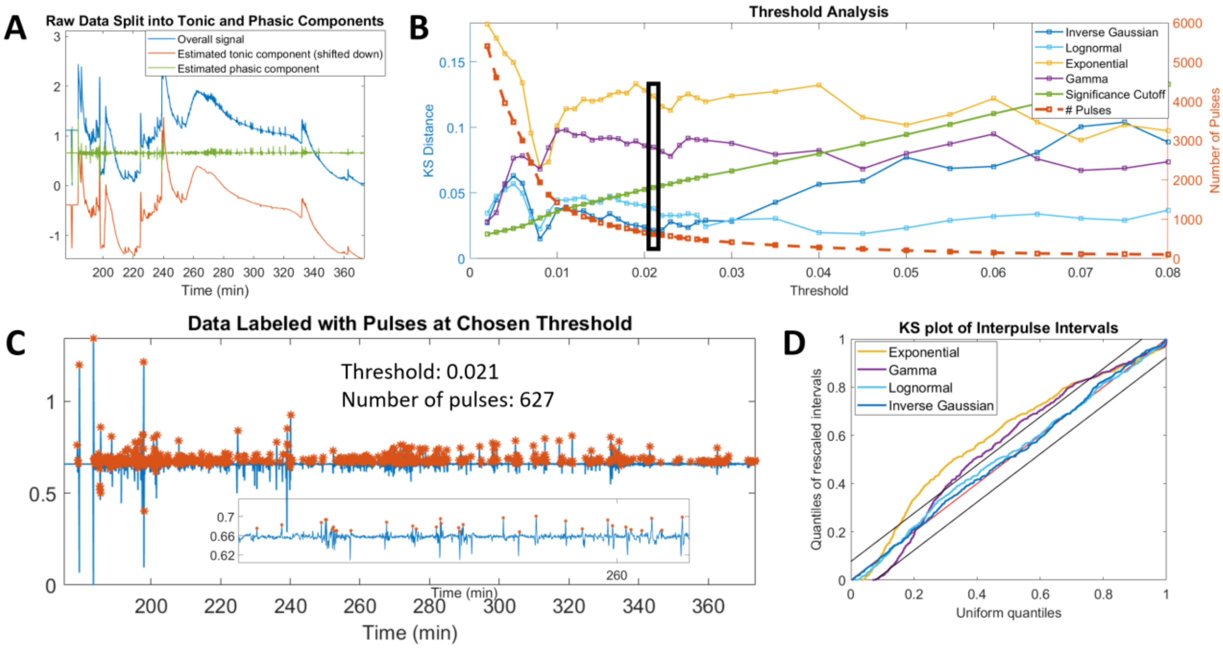


Fig. S18. Results for Subject 10 from the propofol sedation cohort, showing agreement with the trends of the cohort as a whole. (a) Preprocessing of data by splitting into tonic and phasic components, (b) Screening of thresholds with chosen threshold marked with bolded rectangle, (c) Pulses extracted at chosen threshold, (d) Full KS-plot showing goodness-of-fit at chosen threshold

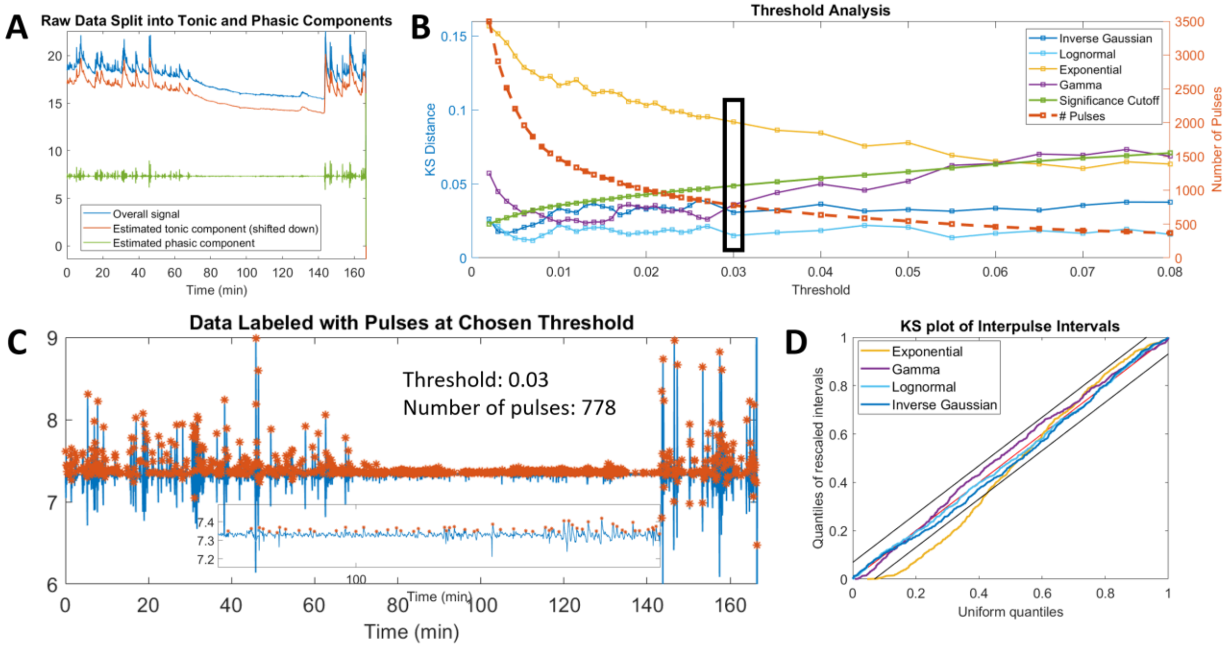


Fig. S19. Results for Subject 11 from the propofol sedation cohort, showing agreement with the trends of the cohort as a whole. (a) Preprocessing of data by splitting into tonic and phasic components, (b) Screening of thresholds with chosen threshold marked with bolded rectangle, (c) Pulses extracted at chosen threshold, (d) Full KS-plot showing goodness-of-fit at chosen threshold

### LEDALAB ALGORITHM RESULTS ON AWAKE AND AT REST COHORT

TABLE S-III
SUMMARY OF KS-DISTANCE RESULTS FOR THE AWAKE AND AT REST COHORT USING THE LEDALAB ALGORITHM

| | Num. pulses | Models under Sig. Cutoff |
|---|---|---|
| **S1** | 727 | IG |
| **S2** | 383 | -- |
| **S3** | 762 | -- |
| **S4** | 1010 | -- |
| **S5** | 566 | -- |
| **S6** | 838 | IG, LogN |
| **S7** | 1250 | -- |
| **S8** | 494 | -- |
| **S9** | 575 | LogN |
| **S10** | 627 | -- |
| **S11** | 778 | -- |

Sig. cutoff = significance cutoff, IG = inverse Gaussian, LogN = lognormal

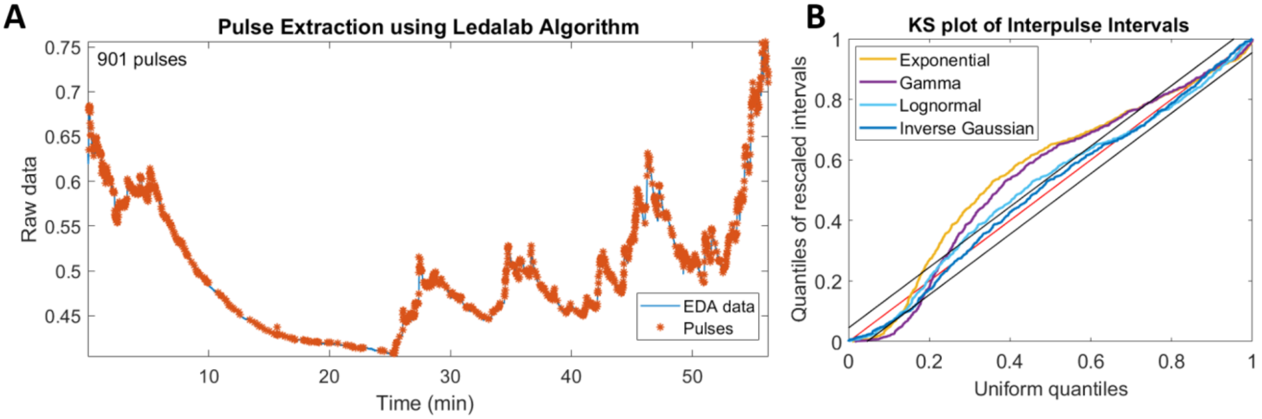


Fig. S20. (a) Pulse selection and (b) goodness-of-fit results using the Ledalab algorithm on S1 from the awake and at rest cohort.

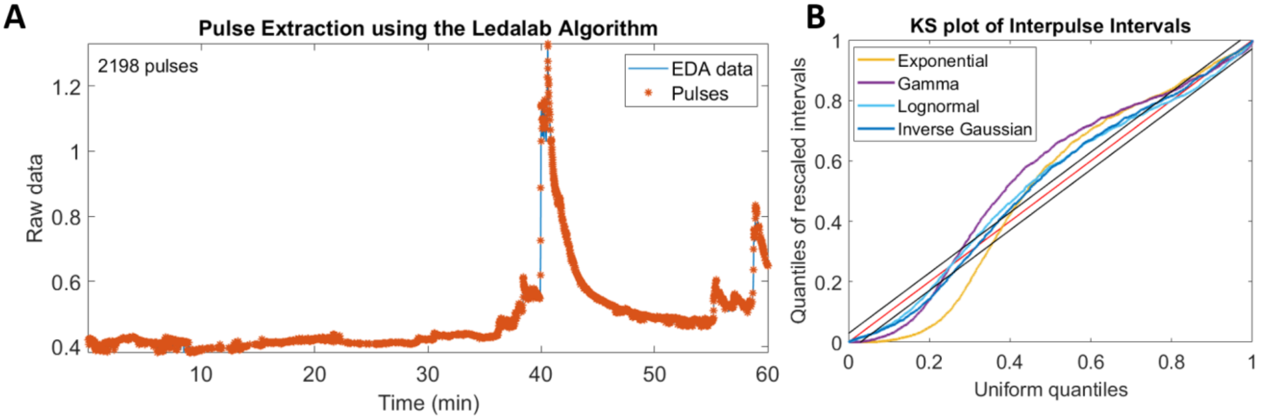


Fig. S21. (a) Pulse selection and (b) goodness-of-fit results using the Ledalab algorithm on S2 from the awake and at rest cohort.

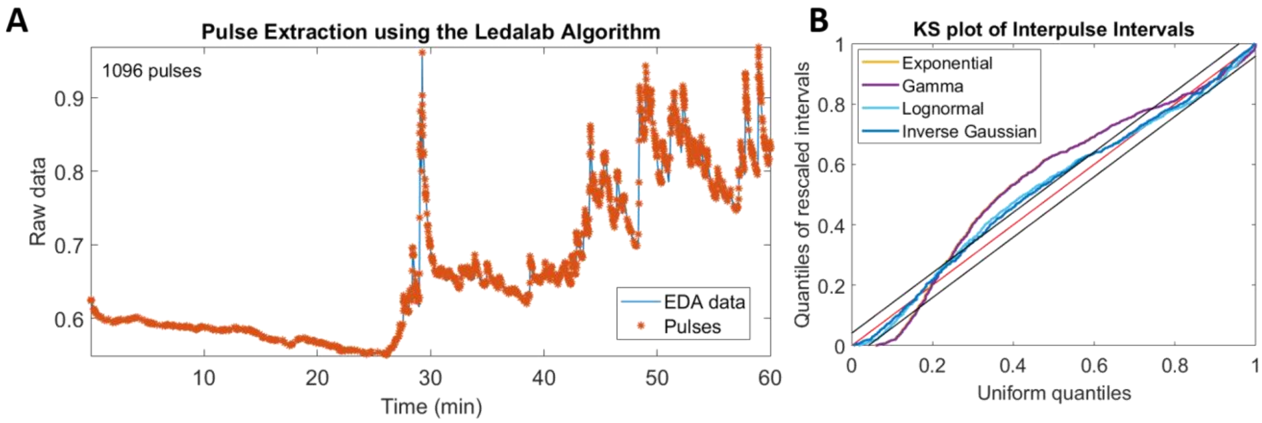


Fig. S22. (a) Pulse selection and (b) goodness-of-fit results using the Ledalab algorithm on S3 from the awake and at rest cohort.

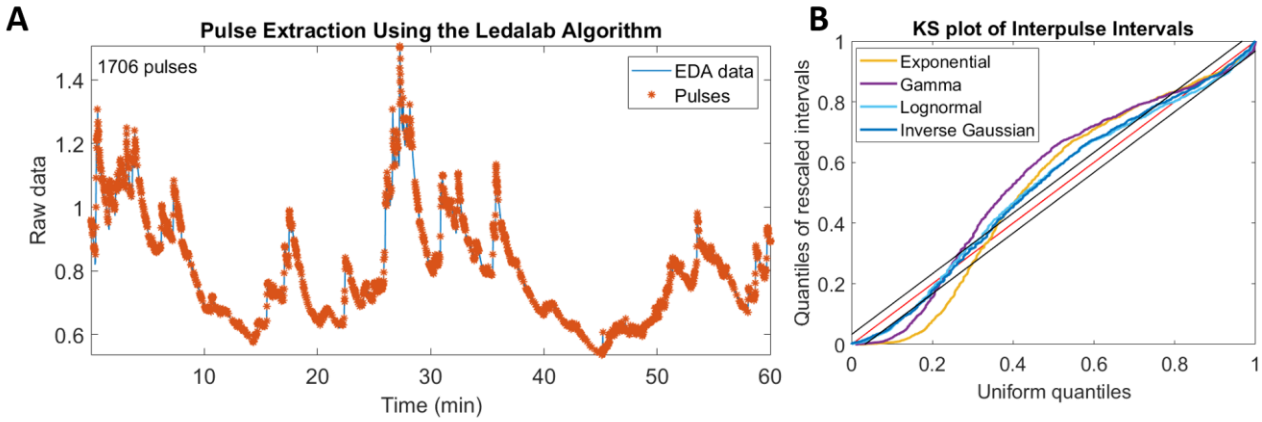


Fig. S23. (a) Pulse selection and (b) goodness-of-fit results using the Ledalab algorithm on S4 from the awake and at rest cohort.

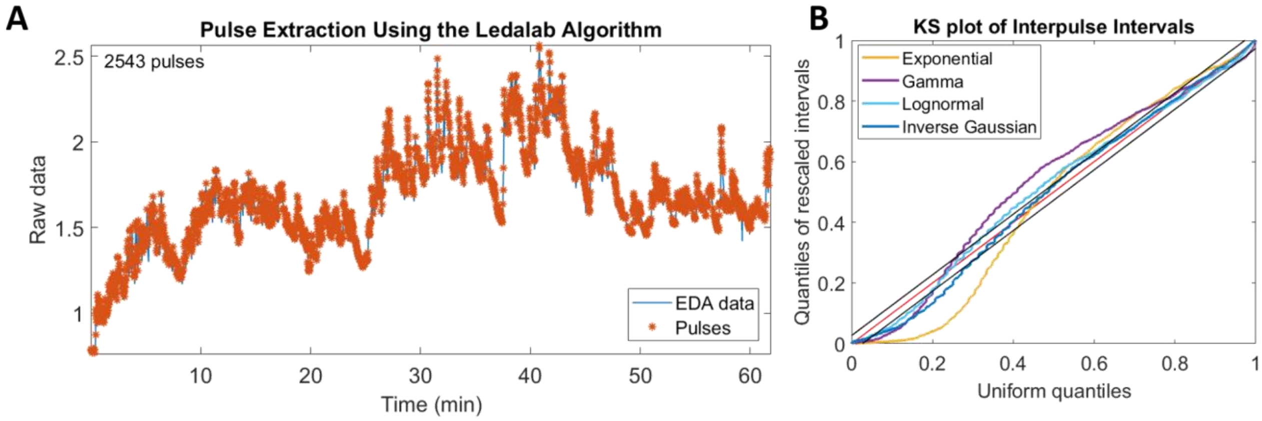


Fig. S24. (a) Pulse selection and (b) goodness-of-fit results using the Ledalab algorithm on S5 from the awake and at rest cohort.

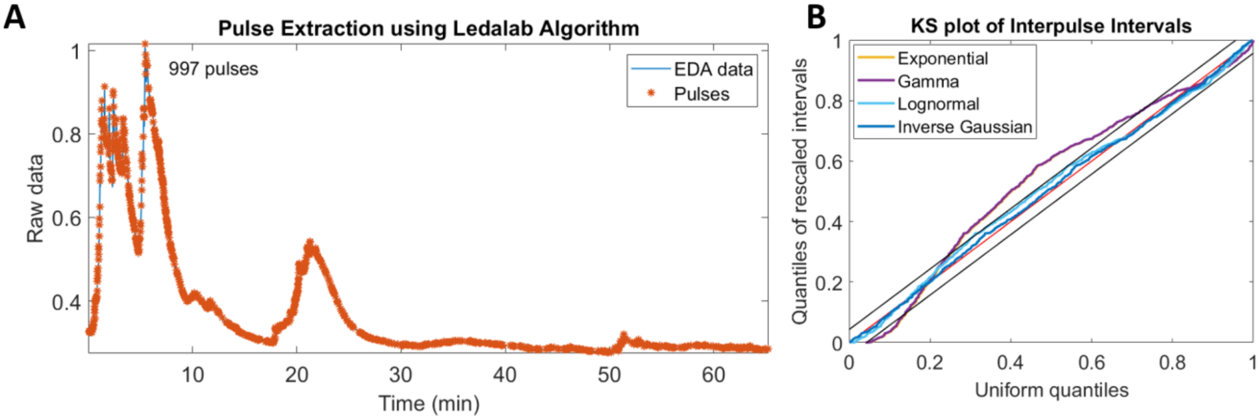


Fig. S25. (a) Pulse selection and (b) goodness-of-fit results using the Ledalab algorithm on S6 from the awake and at rest cohort.

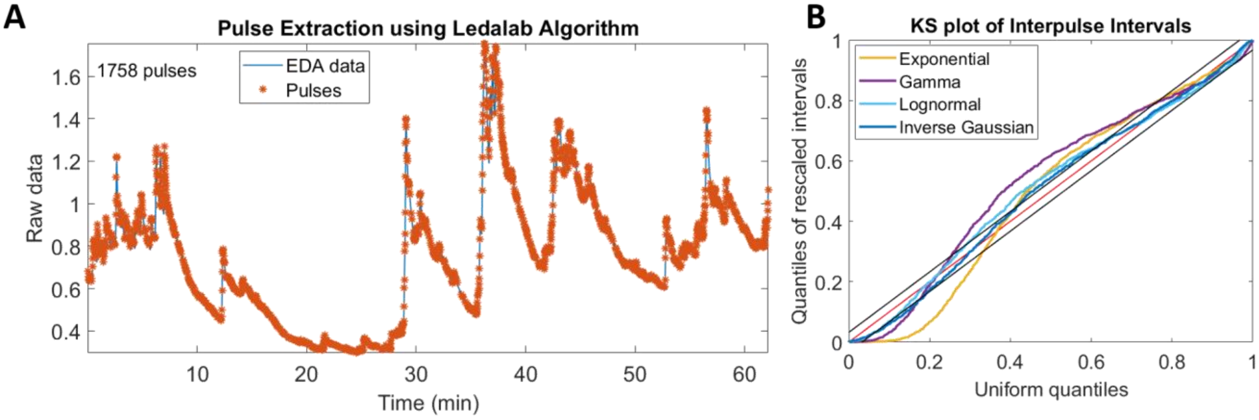


Fig. S26. (a) Pulse selection and (b) goodness-of-fit results using the Ledalab algorithm on S7 from the awake and at rest cohort.

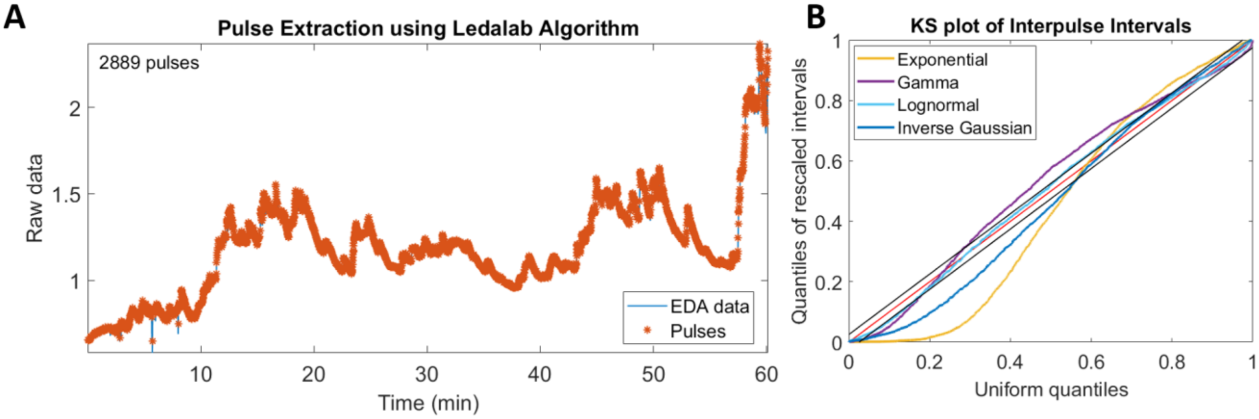


Fig. S27. (a) Pulse selection and (b) goodness-of-fit results using the Ledalab algorithm on S9 from the awake and at rest cohort.

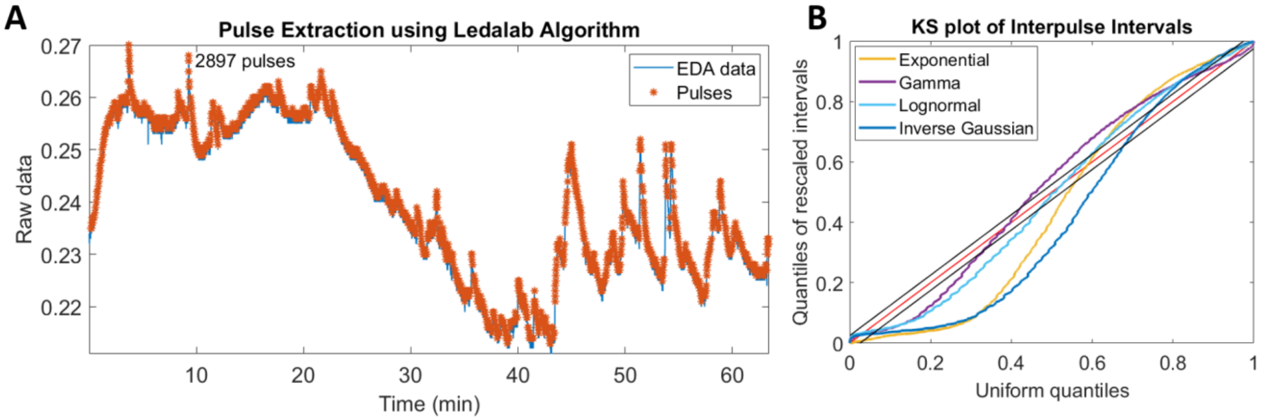


Fig. S28. (a) Pulse selection and (b) goodness-of-fit results using the Ledalab algorithm on S10 from the awake and at rest cohort.

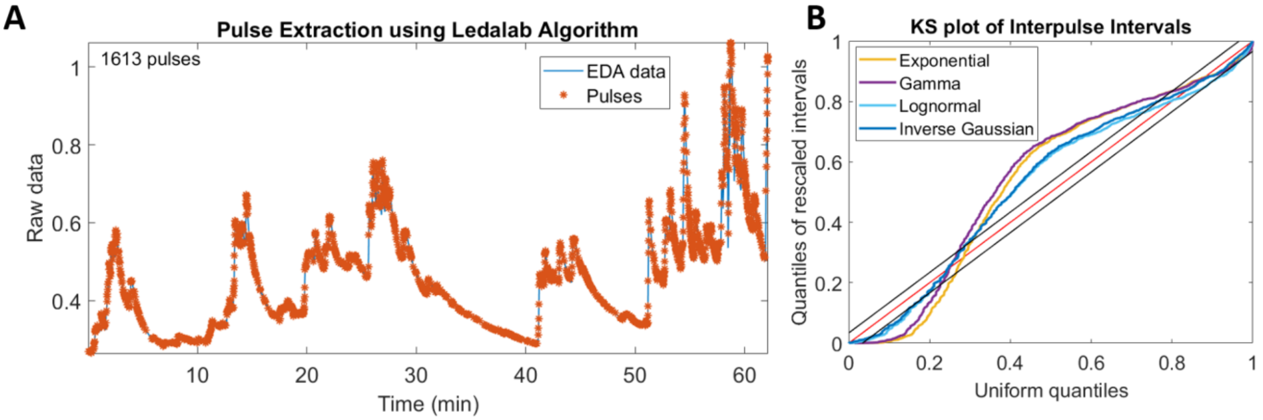


Fig. S29. (a) Pulse selection and (b) goodness-of-fit results using the Ledalab algorithm on S11 from the awake and at rest cohort.
